# Weekly inhaled salmeterol xinafoate, a selective β2 adrenergic receptor agonist drug, therapeutically inhibits cancer growth and prolongs survival *in vivo*

**DOI:** 10.1101/2024.10.15.618606

**Authors:** Xiaofei She, Junxian Ma, Hua Gao

## Abstract

Humans have spent approximately 99.5% of the 2.5 million years of evolutionary history as hunter-gatherers, which helped to evolutionarily mold our minds and bodies to the hunting and gathering lifestyle. Physical activity associated with hunting animals, gathering plants, and evading predators is essential for human survival, evolution, and prosperity. Thus, physical activity (acute stress) is mostly beneficial to the body, whereas psycho-emotional stress (chronic stress) is usually detrimental at the psychological, physiological, and pathological levels. Although physical activity exerts anticancer effects and psycho-emotional stress exhibits cancer-promoting effects, both function through the same pathway involving activation of the sympathetic nervous system, release of catecholamines, and stimulation of β2 adrenergic receptor (β_2_AR) expressed on cancer cells. Here, salmeterol xinafoate was chosen to mimic physical activity. A once-weekly administration of salmeterol xinafoate by the intraperitoneal, inhalation, or transdermal route, dose-dependently extended latency, reduced cancer incidence and growth, and prolonged survival *in vivo*. Remarkably, the best complete response rates were 75%, 30%, and 71.43% for the therapeutic treatment of lung cancer, breast cancer, and melanoma, respectively. Notably, the salmeterol xinafoate doses used in our experiments are comparable to those used in patients. Moreover, in endogenous β_2_AR knockout cancer cells, the exogenous β_2_AR mutant with impaired salmeterol xinafoate binding or downstream signaling prevented the anticancer effects of salmeterol xinafoate *in vivo*, demonstrating the on-target action of salmeterol xinafoate on cancer cells and the requirement of the downstream signaling pathway. Finally, we showed that treatment with salmeterol xinafoate dose-dependently decreased the self-renewal capacity of lung cancer stem cells, breast cancer stem cells, and melanoma stem cells. From the perspective of human evolution into cancer biology, our study reveals that β_2_AR represents an ideal anticancer drug target and that salmeterol xinafoate monotherapy has strong anticancer activity.

## Introduction

For approximately 2.5 million years, humans lived a hunting and gathering lifestyle. Physical activity associated with hunting animals, gathering plants, and evading predators is essential for human survival, evolution, and prosperity (Figure 1A) (*1–6*). Evolutionarily, the human body was adapted to physical activity, and the sympathetic nervous system was selected to respond to physical activity (*1, 7*). Accordingly, the sympathetic nervous system, an integrative system, can directly regulate the function of most organ systems, such as by increasing cardiac output, speeding up the respiratory rate, and boosting blood circulation (*1–3, 7–10*). The sympathetic nervous system also responds to both external dangerous situations (“fight or flight” response) and danger cues from the internal organs of the body to maintain whole-body homeostasis (*8–12*). Sympathetic signaling is achieved through the localized release of catecholamines (mainly norepinephrine) from sympathetic nerve terminals and through the systemic circulation of catecholamines (mainly epinephrine) released from the adrenal gland (Figure 1B) (*8–11, 13*).

**Figure 1.**
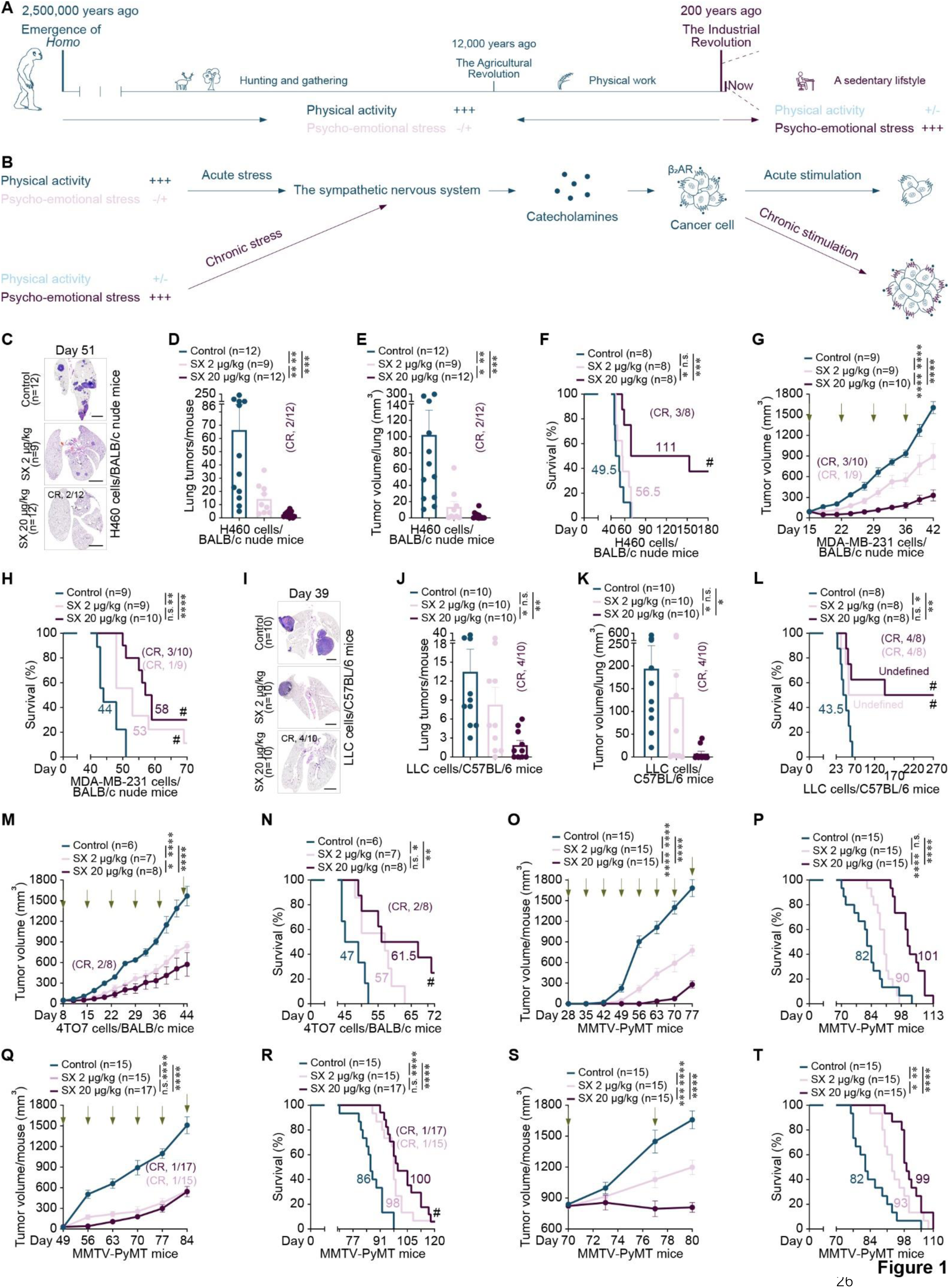
A once-weekly intraperitoneal administration of salmeterol xinafoate therapeutically inhibits the growth of established tumors and prolongs survival of model mice with lung cancer, breast cancer, and melanoma. (A) Brief overview of human evolution. For approximately 2.5 million years, humans lived a hunting and gathering lifestyle. Physical activity associated with hunting animals, gathering plants, and evading predators is essential for human survival, evolution, and prosperity. Evolutionarily, the human body was adapted to physical activity, and the sympathetic nervous system was selected to respond to physical activity. The Industrial Revolution of the last 200 years has replaced human labor with machines and increased the productive potential of humanity. As a result, ever-increasing numbers of people are engaged in office-based vocations, and a sedentary lifestyle is highly prevalent. Therefore, physical activity (acute stress) has been relegated from an absolute necessity of human survival to a choice of maintaining a healthy lifestyle. In addition, psycho-emotional stress (chronic stress), a common factor in modern civilization, has a huge negative impact on our daily lives. Since humans have spent approximately 99.5% of the 2.5 million years of evolutionary history as hunter-gatherers, which helped to evolutionarily mold our minds and bodies to the hunting and gathering lifestyle, and given that human evolution is a gradual process, the body of present-day modern humans is poorly adapted to the sedentary lifestyle. (B) In the context of cancer, accumulated evidence indicates that acute stimulation of β_2_AR expressed on cancer cells [by the evolutionarily selected physical activity (acute stress)–sympathetic nervous system–catecholamines–β_2_AR pathway] can control the initiation, growth, and metastasis of various cancers and improve the overall survival of patients, whereas chronic stimulation of β_2_AR [by the new psycho-emotional stress (chronic stress, a common factor after industrial revolution)–sympathetic nervous system–catecholamines–β_2_AR pathway] exhibits cancer-promoting effects. (**C**-**E**) Representative images of (C) hematoxylin and eosin-stained lung tissue sections, (D) the number of tumors per lung, and (E) the tumor volume per lung at day 51 from BALB/c nude mice xenografted with H460 human lung cancer cells (1 × 10^6^) by intravenous tail vein injection. A once-weekly intraperitoneal administration of salmeterol xinafoate (2 or 20 μg/kg) was initiated from day 21 when the lung tumor was detected (Figure S1A). The experimental endpoint was on day 51 when the first dead mouse was observed in the control group. All lungs were analyzed by serial sectioning the entire lung and scoring tumor nodules. A tumor nodule larger than 100 μm in width was counted and calculated as described in the Materials and Methods. Of the 20 μg/kg treated mice, lung tumors disappeared in 2 of 12 mice. The n-values denote the number of mice per group. SX: Salmeterol xinafoate. CR: Complete response. Scale bars: 2 mm. (**F**) Independent experiment to show the overall survival curves of BALB/c nude mice xenografted with H460 human lung cancer cells (1 × 10^6^) by intravenous tail vein injection. A once-weekly intraperitoneal administration of salmeterol xinafoate (2 or 20 μg/kg) was initiated from day 21 when the lung tumor was detected (Figure S1A). Of the 20 μg/kg-treated mice, 3 of 8 mice did not show signs of illness, and the mice lived for at least 180 days after cancer cell injection until the experimental endpoint at day 180. The n-values denote the number of mice per group. SX: Salmeterol xinafoate. CR: Complete response. # The experimental endpoint on day 180. (**G** and **H**) Primary tumor growth curves (G) and overall survival curves (H) of BALB/c nude mice orthotopically xenografted with MDA-MB-231 human breast cancer cells (1 × 10^6^). A once-weekly intraperitoneal administration of salmeterol xinafoate (2 or 20 μg/kg) was initiated from day 15 when the primary tumor reached approximately 90 mm^3^. The green arrow indicates the time of intraperitoneal injection of salmeterol xinafoate. The primary tumor growth curves were stopped on day 42 when the first primary tumor reached 2000 mm^3^ in the control group (G). Mice were considered to have expired when the primary tumor reached 2000 mm^3^ (H). Of the 20 μg/kg-treated mice, tumors disappeared in 3 of 10 mice, and the 3 mice lived for at least 30, 45, and 57 days, without detectable tumors until the experimental endpoint at day 90. Of the 2 μg/kg-treated mice, tumors disappeared in 1 of 9 mice, and the mice lived for at least 33 days without detectable tumors until the experimental endpoint at day 90. The colored number is the median survival time (H). The n-values denote the number of mice per group. SX: Salmeterol xinafoate. CR: Complete response. # The experimental endpoint on day 90. (**I**-**K**) Representative images of hematoxylin and eosin-stained lung tissue sections (I), the number of tumors per lung (J), and the tumor volume per lung (K) at day 39 from C57BL/6 mice isografted with LLC mouse lung cancer cells (3 × 10^4^) by intravenous tail vein injection. A once-weekly intraperitoneal administration of salmeterol xinafoate (2 or 20 μg/kg) was initiated from day 5 when the lung tumor was detected (Figure S2A). The experimental endpoint was on day 39 when the first dead mouse was observed in the control group. All lungs were analyzed by serial sectioning the entire lung and scoring tumor nodules. A tumor nodule larger than 100 μm in width was counted and calculated as described in the Materials and Methods. Of the 20 μg/kg treated mice, lung tumors disappeared in 4 of 10 mice. The n-values denote the number of mice per group. SX: Salmeterol xinafoate. CR: Complete response. Scale bars: 2 mm. (**L**) Independent experiment to show the overall survival curves of C57BL/6 mice isografted with LLC mouse lung cancer cells (3 × 10^4^) by intravenous tail vein injection. A once-weekly intraperitoneal administration of salmeterol xinafoate (2 or 20 μg/kg) was initiated from day 5 when the lung tumor was detected (Figure S2A). Of the 20 μg/kg-treated mice, 4 of 8 mice did not show signs of illness, and the mice lived for at least 270 days after cancer cell injection until the experimental endpoint at day 270. Of the 2 μg/kg-treated mice, 4 of 8 mice did not show signs of illness, and the mice lived for at least 270 days after cancer cell injection until the experimental endpoint at day 270. The n-values denote the number of mice per group. SX: Salmeterol xinafoate. CR: Complete response. # The experimental endpoint on day 270. (**M** and **N**) Primary tumor growth curves (M) and overall survival curves (N) of BALB/c mice orthotopically isografted with 4TO7 mouse breast cancer cells (1 × 10^6^). A once-weekly intraperitoneal administration of salmeterol xinafoate (2 or 20 μg/kg) was initiated from day 8 when the primary tumor reached approximately 50 mm^3^. The green arrow indicates the time of intraperitoneal injection of salmeterol xinafoate. The primary tumor growth curves were stopped on day 44 when the first primary tumor reached 2000 mm^3^ in the control group (M). Mice were considered to have expired when the primary tumor reached 2000 mm^3^ (N). Of the 20 μg/kg-treated mice, tumors disappeared in 2 of 8 mice, 1 mouse lived for at least 65 days, and 1 mouse lived for at least 59 days without detectable tumors until the experimental endpoint at day 85. The colored number is the median survival time (N). The n-values denote the number of mice per group. SX: Salmeterol xinafoate. CR: Complete response. # The experimental endpoint on day 85. (**O** and **P**) Preventive treatment experiment in the MMTV-PyMT mouse model of spontaneous breast cancer (Figure S2L). Total mammary tumor growth curves (O) and overall survival curves (P) of MMTV-PyMT mice with or without salmeterol xinafoate treatment. A once-weekly intraperitoneal administration of salmeterol xinafoate (2 or 20 μg/kg) was initiated from day 28 when no palpable or visible mammary tumors existed. The green arrow indicates the time of intraperitoneal injection of salmeterol xinafoate. The total mammary tumor growth curves were stopped on day 77 when the first total mammary tumor per mouse reached 2000 mm^3^ in the control group (O). Mice were considered to have expired when the total mammary tumors per mouse reached 2000 mm^3^ (P). The colored number is the median survival time (P). The n-values denote the number of mice per group. SX: Salmeterol xinafoate. (**Q** and **R**) Treatment of early carcinoma in the MMTV-PyMT mouse model of spontaneous breast cancer (Figure S2L). Total mammary tumor growth curves (Q) and overall survival curves (R) of MMTV-PyMT mice with or without salmeterol xinafoate treatment. A once-weekly intraperitoneal administration of salmeterol xinafoate (2 or 20 μg/kg) was initiated from day 49 when palpable or visible mammary tumors existed. The green arrow indicates the time of intraperitoneal injection of salmeterol xinafoate. The total mammary tumor growth curves were stopped on day 84 when the first total mammary tumor per mouse reached 2000 mm^3^ in the control group (Q). Mice were considered to have expired when the total mammary tumors per mouse reached 2000 mm^3^ (R). Of the 20 μg/kg-treated mice, tumors disappeared in 1 of 17 mice, and the mice lived for at least 60 days without detectable tumors until the experimental endpoint at day 180. Of the 2 μg/kg-treated mice, tumors disappeared in 1 of 15 mice, and the mice lived for at least 60 days without detectable tumors until the experimental endpoint at day 180. The colored number is the median survival time (R). The n-values denote the number of mice per group. SX: Salmeterol xinafoate. CR: Complete response. # The experimental endpoint on day 180. (**S** and **T**) Treatment of late carcinoma in the MMTV-PyMT mouse model of spontaneous breast cancer (Figure S2L). Total mammary tumor growth curves (S) and overall survival curves (T) of MMTV-PyMT mice with or without salmeterol xinafoate treatment. A once-weekly intraperitoneal administration of salmeterol xinafoate (2 or 20 μg/kg) was initiated from day 70 when the total mammary tumor reached approximately 800 mm^3^ and lung metastasis developed spontaneously. The green arrow indicates the time of intraperitoneal injection of salmeterol xinafoate. The total mammary tumor growth curves were stopped on day 80 when the first total mammary tumor per mouse reached 2000 mm^3^ in the control group (S). Mice were considered to have expired when the total mammary tumors per mouse reached 2000 mm^3^ (T). The colored number is the median survival time (T). The n-values denote the number of mice per group. SX: Salmeterol xinafoate. Anesthetics were not used in the mouse experiments, except for cancer cells injection, because anesthetics prevented the anticancer effects of salmeterol xinafoate. Mice were fasted for 6 h (H460 and MDA-MB-231) or 12 h (LLC, 4TO7, and MMTV-PyMT mice) before salmeterol xinafoate administration. Four hours (H460 and MDA-MB-231) or 8 h (LLC, 4TO7, and MMTV-PyMT mice) after salmeterol xinafoate administration, food was provided. The data are presented as the mean ± s.e.m. values. *P*-values were determined by unpaired one-way ANOVA with uncorrected Fisher’s least significant difference (LSD) test (control vs. 2 μg/kg or control vs. 20 μg/kg in D, E, J, and K), unpaired two-tailed Student’s t-test with Welch’s correction (2 μg/kg vs. 20 μg/kg in D, E, J, and K), unpaired two-way ANOVA with uncorrected Fisher’s LSD test (G, M, O, Q, and S), or the log-rank test (F, H, L, N, P, R, and T). \**P* < 0.05, \*\**P* < 0.01, \*\*\**P* < 0.001, \*\*\*\**P* < 0.0001, and n.s., not significant.

The Industrial Revolution of the last 200 years has replaced human labor with machines and increased the productive potential of humanity. As a result, ever-increasing numbers of people are engaged in office-based vocations, and a sedentary lifestyle is highly prevalent (Figure 1A). Therefore, physical activity (acute stress) has been relegated from an absolute necessity of human survival to a choice of maintaining a healthy lifestyle (*1, 3, 13–15*). In addition, psycho-emotional stress (chronic stress), a common factor in modern civilization, has a huge negative impact on our daily lives (*16, 17*). Since humans have spent approximately 99.5% of the 2.5 million years of evolutionary history as hunter-gatherers, which helped to evolutionarily mold our minds and bodies to the hunting and gathering lifestyle, and given that human evolution is a gradual process, the body of present-day modern humans is poorly adapted to the sedentary lifestyle (Figure 1A) (*1, 2, 4–6, 13, 14, 16, 18*). Thus, physical activity (acute stress) is mostly beneficial to the body, whereas psycho-emotional stress (chronic stress) is usually detrimental at the psychological, physiological, and pathological levels (*15–17, 19–26*).

Epidemiological studies have demonstrated that physical activity reduces cancer incidence, inhibits cancer growth and metastasis, and lowers the risk of cancer recurrence (*19, 21, 24–26*). Indeed, physical activity-conditioned serum from patients with breast cancer and healthy individuals decreased tumorigenesis in mice preincubated with MCF-7 human breast cancer cells by 50%. Moreover, the blockade of β-adrenergic signaling in MCF-7 cells completely blunted the suppressive effect of the physical activity-conditioned serum on tumorigenesis, indicating that catecholamine was the responsible physical activity factor (*27*). Furthermore, mouse models have shown that physical activity induced a reduction in cancer incidence and growth of more than 60% in different cancer models by enhancing catecholamine levels (*19, 27–30*). By contrast, numerous clinical analyses and an increasing number of *in vivo* studies have shown that psycho-emotional stress promotes the initiation, progression, and metastasis of many types of cancers while also reducing survival (*17, 19, 23, 31–34*). Intriguingly, although physical activity (acute stress) exerts anticancer effects and psycho-emotional stress (chronic stress) exhibits cancer-promoting effects, both function through the same pathway involving activation of the sympathetic nervous system, release of catecholamines from sympathetic nerve terminals (mainly norepinephrine) and the adrenal gland (mainly epinephrine), and stimulation of β2 adrenergic receptor (β_2_AR) expressed on cancer cells (Figure 1B) (*13, 14, 35–50*). However, it is unclear how to mimic or potentiate the evolutionarily selected physical activity–sympathetic nervous system–catecholamines–β_2_AR pathway to use its beneficial anticancer effects in preclinical and clinical cancer therapy (*3, 51–55*).

Here, from the perspective of human evolution into cancer biology, we hypothesized that in the context of cancer, acute stimulation of β_2_AR, which could mimic or potentiate the evolutionarily selected pathway, may reduce cancer incidence, inhibit cancer growth, and prolong survival *in vivo*, since humans have spent approximately 99.5% of the 2.5 million years of evolutionary history as hunter-gatherers, meaning that the body and mind of present-day modern humans remain well adapted to the hunter-gatherer way of life (Figures 1A and 1B) (*56–60*). Thus, salmeterol xinafoate was chosen because it has greatest ability to selectively stimulate β_2_AR out of all β_2_AR agonists, as well as due to its high affinity, a 5- to 20-fold bias toward the activation of Gα_s_-mediated signaling over the stimulation of β-arrestin-mediated effects, weak induction of receptor internalization and β-arrestin translocation, membrane sequestration, long duration (up to 12 h), and non-competition with endogenous catecholamines (Tables S1 and S2) (*61–70*). Moreover, salmeterol xinafoate/corticosteroid combined medication is among some of the most highly prescribed drugs as a bronchodilator for the treatment of asthma and chronic obstructive pulmonary disease; however, mounting evidence indicates that corticosteroids promote cancer growth and metastasis (*71, 72*).

In established tumors, we demonstrated that a once-weekly intraperitoneal injection of salmeterol xinafoate dose-dependently inhibited the growth of established tumors and prolonged survival in immunodeficient mice xenografted with human lung cancer, breast cancer, and melanoma cells. Similar anticancer effects of intraperitoneally injected salmeterol xinafoate were observed in established tumors in immunocompetent mice isografted with mouse lung cancer, breast cancer, and melanoma cells. This is likely because the salmeterol xinafoate binding sites of human and mouse β_2_AR are identical, and the protein sequence identity between human and mouse β_2_AR is approximately 87%. Furthermore, in the MMTV-PyMT mouse model of spontaneous breast cancer, before primary tumor onset, a once-weekly intraperitoneal injection of salmeterol xinafoate dose-dependently extended latency, reduced cancer incidence and growth, and prolonged survival, while after primary tumor onset, a once-weekly intraperitoneal injection of salmeterol xinafoate dose-dependently inhibited the growth of established tumors and prolonged survival. Intratumoral salmeterol xinafoate was also detected by mass spectrometry. Fascinatingly, both a once-weekly inhalation of salmeterol xinafoate extended latency, inhibited lung and breast cancer incidence and growth, and prolonged survival *in vivo*; and a once-weekly transdermal administration of salmeterol xinafoate inhibited melanoma growth and prolonged survival *in vivo*, as effectively as a once-weekly intraperitoneal injection of salmeterol xinafoate. Moreover, in endogenous β_2_AR knockout cancer cells, the exogenous β_2_AR mutant with impaired salmeterol xinafoate binding or with impaired downstream signaling prevented the anticancer effects of salmeterol xinafoate *in vivo*, demonstrating the on-target action of salmeterol xinafoate on cancer cells, which is dependent on the downstream signaling pathway. Finally, we showed that treatment with salmeterol xinafoate dose-dependently decreased the self-renewal capacity of lung cancer stem cells, breast cancer stem cells, and melanoma stem cells. In addition, it is worth noting that the salmeterol xinafoate doses used in our experiments are comparable to those used in patients with asthma or chronic obstructive pulmonary disease (Tables S4 and S5).

## Results

### Weekly intraperitoneal injection of salmeterol xinafoate therapeutically inhibits the growth of established tumors and prolongs survival *in vivo*

The evolutionarily selected physical activity–sympathetic nervous system– catecholamines–β_2_AR pathway has numerous beneficial effects on many types of cancers (Figures 1A and 1B). To mimic or potentiate the beneficial anticancer effects, salmeterol xinafoate was chosen because it has greatest ability to selectively stimulate β_2_AR out of all β_2_AR agonists, as well as due to its high affinity, a 5- to 20-fold bias toward the activation of Gα_s_-mediated signaling over the stimulation of β-arrestin-mediated effects, weak induction of receptor internalization and β-arrestin translocation, membrane sequestration, long duration (up to 12 h), and non-competition with endogenous catecholamines (Tables S1 and S2) (*61–70*).

For established tumors, a once-weekly intraperitoneal injection of salmeterol xinafoate dose-dependently inhibited the growth of established tumors and prolonged survival of immunodeficient mice xenografted with human lung cancer (Figures 1C-1F and S1A), breast cancer (Figures 1G, 1H, S1B, and S1C), and melanoma cells (Figures S1D-S1G). Unexpectedly, for lung cancer, 3 of the 8 mice of the 20 μg/kg-treated group did not show signs of illness, and the mice lived for at least 180 days after cancer cell injection until the experimental endpoint at day 180 (Figure 1F). For breast cancer, of the 20 μg/kg-treated mice, tumors disappeared in 3 of 10 mice, and the 3 mice lived for at least 30, 45, and 57 days, without detectable tumors until the experimental endpoint at day 90. Of the 2 μg/kg-treated mice, tumors disappeared in 1 of 9 mice, and the mice lived for at least 33 days without detectable tumors until the experimental endpoint at day 90 (Figures 1G, 1H, S1B, and S1C).

The salmeterol xinafoate binding sites of human and mouse β_2_AR are identical, and the protein sequence identity between humans and mouse β_2_AR is approximately 87% (*62, 64*). Therefore, we speculated that acute stimulation of β_2_AR exerts anticancer effects on mouse cancer cells as effectively as on human cancer cells. Indeed, for established tumors, a once-weekly intraperitoneal injection of salmeterol xinafoate dose-dependently inhibited the growth of established tumors and prolonged survival of immunocompetent mice isografted with mouse lung cancer (Figures 1I-1L and S2A), breast cancer (Figures 1M, 1N, and S2B-S2G), and melanoma cells (Figures S2H-S2K). For lung cancer, 4 of the 8 mice of the 20 μg/kg-treated group did not show signs of illness and lived for at least 270 days after cancer cell injection until the experimental endpoint at day 270. Of the 2 μg/kg-treated mice, 4 of 8 mice did not show signs of illness and lived for at least 270 days after cancer cell injection until the experimental endpoint at day 270 (Figure 1L). For 4TO7 breast cancer, of the 20 μg/kg-treated mice, tumors disappeared in 2 of 8 mice, 1 mouse lived for at least 65 days, and 1 mouse lived for at least 59 days without detectable tumors until the experimental endpoint at day 85 (Figures 1M, 1N, and S2B).

Moreover, in the MMTV-PyMT mouse model of spontaneous breast cancer (Figure S2L), we found that before primary tumor onset (28 days after birth), a once-weekly intraperitoneal injection of salmeterol xinafoate dose-dependently extended latency (Figure S2M), reduced cancer incidence and growth (Figures 1O and S2N-S2P), and prolonged survival (Figure 1P). For established tumors in MMTV-PyMT mice (49 or 70 days after birth), a once-weekly intraperitoneal injection of salmeterol xinafoate dose-dependently inhibited the growth of established tumors and prolonged survival (Figures 1Q-1T and S2Q-S2W). Intratumoral salmeterol xinafoate was also detected using mass spectrometry (Table S3).

It should be noted that the salmeterol xinafoate doses used in mice are less than those used in patients with asthma or chronic obstructive pulmonary disease (Table S4). These results also suggest that the anticancer function of salmeterol xinafoate is independent of the immune system.

### Weekly inhalational or transdermal administration of salmeterol xinafoate therapeutically inhibits the growth of established tumors and prolongs survival *in vivo*

Inhaled medications are ideal for treating respiratory diseases because of the direct interactions of drug and pharmacological targets (*73–76*). Furthermore, inhaled salmeterol xinafoate/corticosteroid combined medication is among some of the most commonly prescribed drugs used to treat asthma and chronic obstructive pulmonary disease; however, mounting evidence indicates that corticosteroids promote cancer growth and metastasis (*71, 72*). Thus, we next investigated whether inhaled salmeterol xinafoate as monotherapy exerts anticancer effects on lung cancer *in vivo*. Intriguingly, a once-weekly inhalation of salmeterol xinafoate dose-dependently inhibited the growth of established tumors and prolonged survival in immunodeficient mice xenografted with H460 human lung cancer cells (Figures 2A-2D and S1A). Of the 6 μg/mouse-treated mice, 4 of 8 mice did not show signs of illness and lived for at least 180 days after cancer cell injection until the experimental endpoint at day 180. Of the 2 μg/mouse-treated mice, 1 of 8 mice did not show signs of illness and lived for at least 180 days after cancer cell injection until the experimental endpoint at day 180 (Figure 2D).

**Figure 2.**
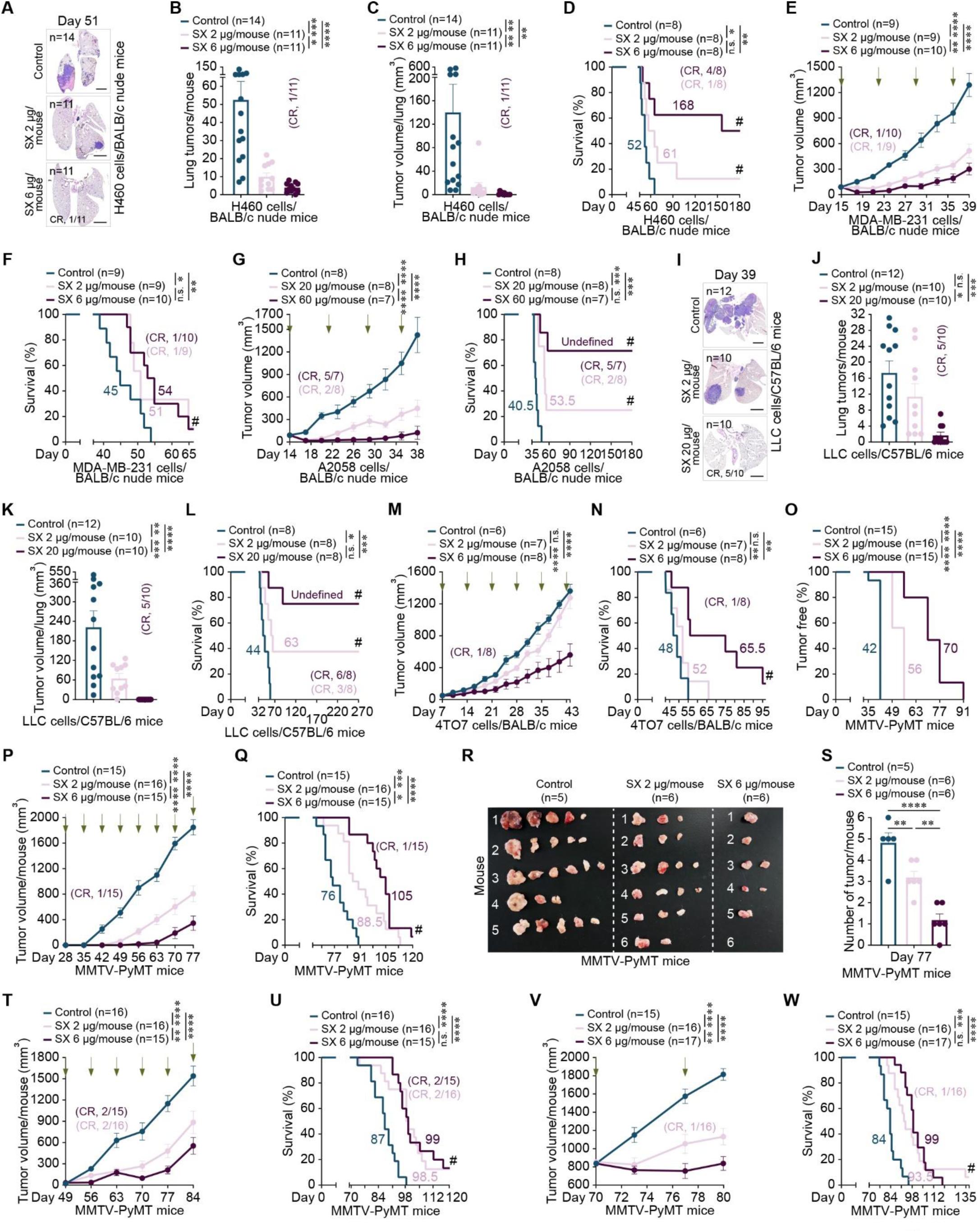
A once-weekly inhalational or transdermal administration of salmeterol xinafoate therapeutically inhibits the growth of established tumors and prolongs survival of model mice with lung cancer, breast cancer, and melanoma. (**A**-**C**) Representative images of hematoxylin and eosin-stained lung tissue sections (A), the number of tumors per lung (B), and the tumor volume per lung (C) at day 51 from BALB/c nude mice xenografted with H460 human lung cancer cells (1 × 10^6^) by intravenous tail vein injection. A once-weekly inhalational administration of salmeterol xinafoate (2 or 6 μg/mouse) was initiated from day 21 when the lung tumor was detected (Figure S1A). The experimental endpoint was on day 51 when the first dead mouse was observed in the control group. All lungs were analyzed by serial sectioning the entire lung and scoring tumor nodules. A tumor nodule larger than 100 μm in width was counted and calculated as described in the Materials and Methods. Of the 6 μg/mouse treated mice, lung tumors disappeared in 1 of 11 mice. The n-values denote the number of mice per group. SX: Salmeterol xinafoate. CR: Complete response. Scale bars: 2 mm. (**D**) Independent experiment to show the overall survival curves of BALB/c nude mice xenografted with H460 human lung cancer cells (1 × 10^6^) by intravenous tail vein injection. A once-weekly inhalational administration of salmeterol xinafoate (2 or 6 μg/mouse) was initiated from day 21 when the lung tumor was detected (Figure S1A). Of the 6 μg/mouse-treated mice, 4 of 8 mice did not show signs of illness, and the mice lived for at least 180 days after cancer cell injection until the experimental endpoint at day 180. Of the 2 μg/mouse-treated mice, 1 of 8 mice did not show signs of illness, and the mice lived for at least 180 days after cancer cell injection until the experimental endpoint at day 180. The n-values denote the number of mice per group. SX: Salmeterol xinafoate. CR: Complete response. # The experimental endpoint on day 180. (**E** and **F**) Primary tumor growth curves (E) and overall survival curves (F) of BALB/c nude mice orthotopically xenografted with MDA-MB-231 human breast cancer cells (1 × 10^6^). A once-weekly inhalational administration of salmeterol xinafoate (2 or 6 μg/mouse) was initiated from day 15 when the primary tumor reached approximately 90 mm^3^. The green arrow indicates the time of inhalational administration of salmeterol xinafoate. The primary tumor growth curves were stopped on day 39 when the first primary tumor reached 2000 mm^3^ in the control group (E). Mice were considered to have expired when the primary tumor reached 2000 mm^3^ (F). Of the 6 μg/mouse-treated mice, tumors disappeared in 1 of 10 mice, and the mice lived for at least 63 days without detectable tumors until the experimental endpoint at day 90. Of the 2 μg/mouse-treated mice, tumors disappeared in 1 of 9 mice, and the mice lived for at least 60 days without detectable tumors until the experimental endpoint at day 90. The colored number is the median survival time (F). The n-values denote the number of mice per group. SX: Salmeterol xinafoate. CR: Complete response. # The experimental endpoint on day 90. (**G** and **H**) Primary tumor growth curves (G) and overall survival curves (H) of BALB/c nude mice orthotopically xenografted with A2058 human melanoma cells (1 × 10^6^). A once-weekly transdermal administration of salmeterol xinafoate (20 or 60 μg/mouse) was initiated from day 14 when the primary tumor reached approximately 100 mm^3^. The green arrow indicates the time of transdermal administration of salmeterol xinafoate. The primary tumor growth curves were stopped on day 38 when the first primary tumor reached 2000 mm^3^ in the control group (G). Mice were considered to have expired when the primary tumor reached 2000 mm^3^ (H). Of the 60 μg/mouse-treated mice, melanoma disappeared in 5 of 7 mice, 4 mice lived for at least 157 days, and 1 mouse lived for at least 154 days without detectable tumors until the experimental endpoint at day 180. Of the 20 μg/mouse-treated mice, melanoma disappeared in 2 of 8 mice, 1 mouse lived for at least 154 days, and 1 mouse lived for at least 148 days without detectable tumors until the experimental endpoint at day 180. The colored number is the median survival time (H). The n-values denote the number of mice per group. SX: Salmeterol xinafoate. CR: Complete response. # The experimental endpoint on day 180. (**I**-**K**) Representative images of hematoxylin and eosin-stained lung tissue sections (I), the number of tumors per lung (J), and the tumor volume per lung (K) at day 39 from C57BL/6 mice isografted with LLC mouse lung cancer cells (3 × 10^4^) by intravenous tail vein injection. A once-weekly inhalational administration of salmeterol xinafoate (2 or 20 μg/mouse) was initiated from day 5 when the lung tumor was detected (Figure S2A). The experimental endpoint was on day 39 when the first dead mouse was observed in the control group. All lungs were analyzed by serial sectioning the entire lung and scoring tumor nodules. A tumor nodule larger than 100 μm in width was counted and calculated as described in the Materials and Methods. Of the 20 μg/mouse treated mice, lung tumors disappeared in 5 of 10 mice. The n-values denote the number of mice per group. SX: Salmeterol xinafoate. CR: Complete response. Scale bars: 2 mm. (**L**) Independent experiment to show the overall survival curves of C57BL/6 mice isografted with LLC mouse lung cancer cells (3 × 10^4^) by intravenous tail vein injection. A once-weekly inhalational administration of salmeterol xinafoate (2 or 20 μg/mouse) was initiated from day 5 when the lung tumor was detected (Figure S2A). Of the 20 μg/mouse-treated mice, 6 of 8 mice did not show signs of illness, and the mice lived for at least 270 days after cancer cell injection until the experimental endpoint at day 270. Of the 2 μg/mouse-treated mice, 3 of 8 mice did not show signs of illness, and the mice lived for at least 270 days after cancer cell injection until the experimental endpoint at day 270. The n-values denote the number of mice per group. SX: Salmeterol xinafoate. CR: Complete response. # The experimental endpoint on day 270. (**M** and **N**) Primary tumor growth curves (M) and overall survival curves (N) of BALB/c mice orthotopically isografted with 4TO7 mouse breast cancer cells (1 × 10^6^). A once-weekly inhalational administration of salmeterol xinafoate (2 or 6 μg/mouse) was initiated from day 7 when the primary tumor reached approximately 50 mm^3^. The green arrow indicates the time of inhalational administration of salmeterol xinafoate. The primary tumor growth curves were stopped on day 43 when the first primary tumor reached 2000 mm^3^ in the control group (M). Mice were considered to have expired when the primary tumor reached 2000 mm^3^ (N). Of the 6 μg/mouse-treated mice, tumors disappeared in 1 of 8 mice, and the mice lived for at least 44 days without detectable tumors until the experimental endpoint at day 96. The colored number is the median survival time (N). The n-values denote the number of mice per group. SX: Salmeterol xinafoate. CR: Complete response. # The experimental endpoint on day 96. (**O**-**Q**) Preventive treatment experiment in the MMTV-PyMT mouse model of spontaneous breast cancer (Figure S2L). Latency of spontaneous mammary tumors (O), total mammary tumor growth curves (P), and overall survival curves (Q) of MMTV-PyMT mice with or without salmeterol xinafoate treatment. A once-weekly inhalational administration of salmeterol xinafoate (2 or 6 μg/mouse) was initiated from day 28 when no palpable or visible mammary tumors existed. The colored number is the median latency time (O). The green arrow indicates the time of inhalational administration of salmeterol xinafoate (P). The total mammary tumor growth curves were stopped on day 77 when the first total mammary tumor per mouse reached 2000 mm^3^ in the control group (P). Mice were considered to have expired when the total mammary tumors per mouse reached 2000 mm^3^ (Q). Of the 6 μg/mouse-treated mice, tumors disappeared in 1 of 15 mice, and the mice lived for at least 61 days without detectable tumors until the experimental endpoint at day 180. The colored number is the median survival time (Q). The n-values denote the number of mice per group. SX: Salmeterol xinafoate. CR: Complete response. # The experimental endpoint on day 180. (**R** and **S**) Independent preventive treatment experiment in the MMTV-PyMT mouse model of spontaneous breast cancer showing the total mammary tumors (R) and the number of total mammary tumors per mouse (**S**) with or without salmeterol xinafoate treatment at day 77. A once-weekly inhalational administration of salmeterol xinafoate (2 or 6 μg/mouse) was initiated from day 28 when no palpable or visible mammary tumors existed. The n-values denote the number of mice per group. SX: Salmeterol xinafoate. (**T** and **U**) Treatment of early carcinoma in the MMTV-PyMT mouse model of spontaneous breast cancer (Figure S2L). Total mammary tumor growth curves (T) and overall survival curves (U) of MMTV-PyMT mice with or without salmeterol xinafoate treatment. A once-weekly inhalational administration of salmeterol xinafoate (2 or 6 μg/mouse) was initiated from day 49 when palpable or visible mammary tumors existed. The green arrow indicates the time of inhalational administration of salmeterol xinafoate. The total mammary tumor growth curves were stopped on day 84 when the first total mammary tumor per mouse reached 2000 mm^3^ in the control group (T). Mice were considered to have expired when the total mammary tumors per mouse reached 2000 mm^3^ (U). Of the 6 μg/mouse-treated mice, tumors disappeared in 2 of 15 mice, 1 mouse lived for at least 96 days, and 1 mouse lived for at least 82 days without detectable tumors until the experimental endpoint at day 180. Of the 2 μg/mouse-treated mice, tumors disappeared in 2 of 16 mice, and the mice lived for at least 68 days without detectable tumors until the experimental endpoint at day 180. The colored number is the median survival time (U). The n-values denote the number of mice per group. SX: Salmeterol xinafoate. CR: Complete response. # The experimental endpoint on day 180. (**V** and **W**) Treatment of late carcinoma in the MMTV-PyMT mouse model of spontaneous breast cancer (Figure S2L). Total mammary tumor growth curves (V) and overall survival curves (W) of MMTV-PyMT mice with or without salmeterol xinafoate treatment. A once-weekly inhalational administration of salmeterol xinafoate (2 or 6 μg/mouse) was initiated from day 70 when the total mammary tumor reached approximately 800 mm^3^ and lung metastasis developed spontaneously. The green arrow indicates the time of inhalational administration of salmeterol xinafoate. The total mammary tumor growth curves were stopped on day 80 when the first total mammary tumor per mouse reached 2000 mm^3^ in the control group (V). Mice were considered to have expired when the total mammary tumors per mouse reached 2000 mm^3^ (W). Of the 2 μg/mouse-treated mice, tumors disappeared in 1 of 16 mice, and the mice lived for at least 54 days without detectable tumors until the experimental endpoint at day 180. The colored number is the median survival time (W). The n-values denote the number of mice per group. SX: Salmeterol xinafoate. CR: Complete response. # The experimental endpoint on day 180. Anesthetics were not used in the mouse experiments, except for cancer cells injection, because anesthetics prevented the anticancer effects of salmeterol xinafoate. Mice were fasted for 6 h (H460, MDA-MB-231, and A2058) or 12 h (LLC, 4TO7, and MMTV-PyMT mice) before salmeterol xinafoate administration. Four hours (H460, MDA-MB-231, and A2058) or 8 h (LLC, 4TO7, and MMTV-PyMT mice) after salmeterol xinafoate administration, food was provided. The data are presented as the mean ± s.e.m. values. *P*-values were determined by unpaired one-way ANOVA with uncorrected Fisher’s LSD test (control vs. 2 μg/mouse or control vs. 20 μg/mouse in B, C, J, and K), unpaired two-tailed Student’s t-test with Welch’s correction (2 μg/kg vs. 20 μg/kg in B, C, J, and K), unpaired two-way ANOVA with uncorrected Fisher’s LSD test (E, G, M, P, T, and V), or the log-rank test (D, F, H, L, N, O, Q, U, and W). \**P* < 0.05, \*\**P* < 0.01, \*\*\**P* < 0.001, \*\*\*\**P* < 0.0001, and n.s., not significant.

Considerable evidence suggests that the maximum plasma concentration is 0.1-0.2 μg/L 5 min after inhalation of a single 50 μg dose of salmeterol xinafoate (*75, 77, 78*). We further investigated whether inhaled salmeterol xinafoate exerts anticancer effects on human breast cancer *in vivo* because inhaled salmeterol xinafoate can be detected in plasma. Fascinatingly, a once-weekly inhalation of salmeterol xinafoate dose-dependently inhibited the growth of established tumors and prolonged survival in immunodeficient mice orthotopically xenografted with MDA-MB-231 human breast cancer cells (Figures 2E, 2F, S3A, and S3B). Of the 6 μg/mouse-treated mice, tumors disappeared in 1 of 10 mice and the mice lived for at least 63 days without detectable tumors until the experimental endpoint on day 90. Of the 2 μg/mouse-treated mice, tumors disappeared in 1 of 9 mice and the mice lived for at least 60 days without detectable tumors until the experimental endpoint on day 90.

Remarkably, we observed superior anticancer effects of transdermal administered salmeterol xinafoate on established tumors in immunodeficient mice orthotopically xenografted with human melanoma cells (Figures 2G, 2H, S3C, and S3D). Of the 60 μg/mouse-treated mice, melanoma disappeared in 5 of 7 mice, 4 mice lived for at least 157 days, and 1 mouse lived for at least 154 days without detectable tumors until the experimental endpoint at day 180. Of the 20 μg/mouse-treated mice, melanoma disappeared in 2 of 8 mice, 1 mouse lived for at least 154 days, and 1 mouse lived for at least 148 days without detectable tumors until the experimental endpoint at day 180 (Figures 2G, 2H, S3C, and S3D).

Similar anticancer effects of inhaled salmeterol xinafoate were observed in established tumors in mice isografted with mouse lung (Figures 2I-2L and S2A) and breast cancer cells (Figures 2M, 2N, and S4A-S4F). For lung cancer, of the 20 μg/mouse-treated mice, 6 of 8 mice did not show signs of illness and lived for at least 270 days after cancer cell injection until the experimental endpoint at day 270; of the 2 μg/kg-treated mice, 3 of 8 mice did not show signs of illness and lived for at least 270 days after cancer cell injection until the experimental endpoint at day 270 (Figure 2L). For 4TO7 breast cancer, of the 6 μg/mouse-treated mice, tumors disappeared in 1 of 8 mice, and the mice lived for at least 44 days without detectable tumors until the experimental endpoint at day 96 (Figures 2M, 2N, and S4A). Furthermore, in the MMTV-PyMT mouse model of spontaneous breast cancer (Figure S2L), before primary tumor onset (28 days after birth), a once-weekly inhalation of salmeterol xinafoate dose-dependently extended latency, reduced cancer incidence and growth, and prolonged survival (Figures 2O-2S and S4G-S4J). For established tumors (49 or 70 days after birth), a once-weekly inhalation of salmeterol xinafoate dose-dependently inhibited the growth of established tumors and prolonged survival in MMTV-PyMT mice (Figures 2T-2W and S4K-S4Q). In addition, we observed the anticancer effects of a once-weekly transdermal administration of salmeterol xinafoate in established tumors in mice orthotopically isografted with mouse melanoma cells (Figures S4R-S4U). Of the 20 μg/mouse-treated mice, melanoma disappeared in 1 of 10 mice, and the mice lived for at least 116 days without detectable tumors until the experimental endpoint at day 140 (Figures S4R, S4T, and S4U).

It should be noted that the salmeterol xinafoate doses used in mice are similar to those used in patients with asthma or chronic obstructive pulmonary disease (Table S5). These results indicate that both a once-weekly inhalation of salmeterol xinafoate exerts anticancer effects on lung or breast cancer and a once-weekly transdermal administration of salmeterol xinafoate exerts anticancer effects on melanoma *in vivo*, as effectively as a once-weekly intraperitoneal injection of salmeterol xinafoate.

### Salmeterol xinafoate exerts anticancer effects through β_2_AR

To confirm that the anticancer effects of salmeterol xinafoate *in vivo* function directly through the β_2_AR, both the mutant human β_2_AR with F193W, F194V, S203T, S204T, H296K, R304P, K305D, and Y308F mutations (SBD β_2_AR) and the mutant mouse β_2_AR with F193W, F194V, S203T, S204T, H296K, K305D, and Y308F mutations (SBD β_2_AR), which do not bind to salmeterol xinafoate (*62, 64*), were applied to tumorigenesis experiments in mice. In endogenous β_2_AR knockout cancer cells, the exogenous β_2_AR mutant with impaired salmeterol xinafoate binding completely prevented the anticancer effects of salmeterol xinafoate administered through inhalation (lung cancer) or intraperitoneal injection (breast cancer) in immunodeficient mice xenografted with human lung (Figures 3A-3F and S5) or breast cancer cells (Figures 3G-3J and S6) and in immunocompetent mice isografted with mouse lung (6 μg/mouse, Figures 3K-3P and S7) or breast cancer cells (2 μg/kg, Figures 3Q, 3R, and S8), demonstrating the on-target action of salmeterol xinafoate on cancer cells.

**Figure 3.**
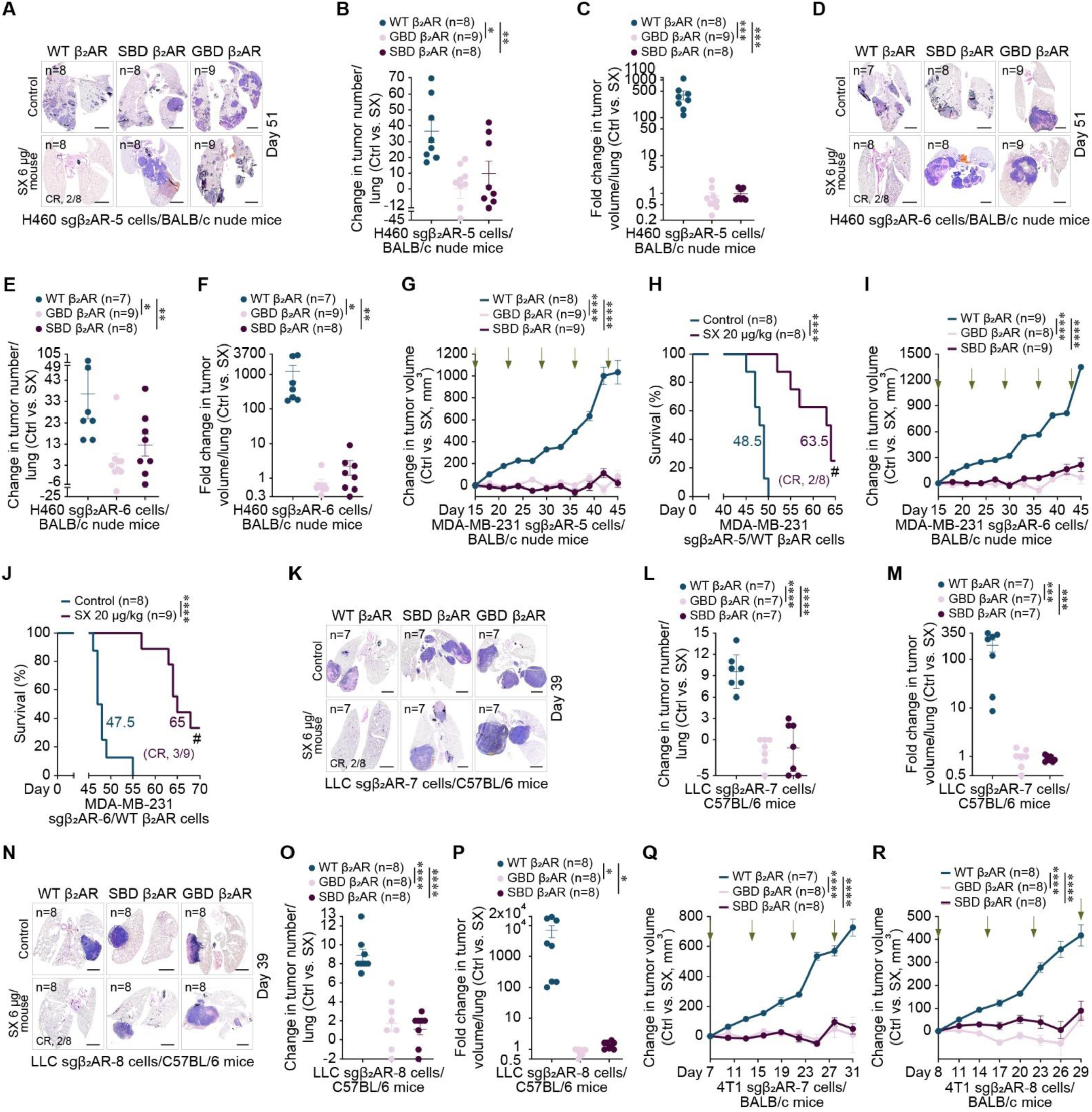
Salmeterol xinafoate exerts anticancer effects *in vivo* through β_2_AR expressed on cancer cells and is dependent on the downstream signaling pathway. (**A**-**C**) Representative images of hematoxylin and eosin-stained lung tissue sections (A), change in the tumor number per lung between the control and salmeterol xinafoate treatment groups (B), and the fold change in tumor volume per lung between the control and salmeterol xinafoate treatment groups (C) on day 51 from BALB/c nude mice xenografted with the indicated H460 human lung cancer cells (1 × 10^6^) by intravenous tail vein injection. In endogenous β_2_AR knockout H460 lung cancer cells (sgβ_2_AR-5), wild-type human β_2_AR (WT β_2_AR), salmeterol xinafoate-binding-deficient human β_2_AR (F193W, F194V, S203T, S204T, H296K, R304P, K305D, and Y308F mutations, SBD β_2_AR), or G-protein-binding-deficient human β_2_AR (R131A and F139A mutations, GBD β_2_AR) was overexpressed as indicated. A once-weekly inhalational administration of salmeterol xinafoate (6 μg/mouse) was initiated from day 21 when the lung tumor was detected (Figure S1A). The experimental endpoint was on day 51 when the first dead mouse was observed in the control group. All lungs were analyzed by serial sectioning the entire lung and scoring the tumor nodules. A tumor nodule larger than 100 μm in width was counted and calculated as described in the Materials and Methods. Of the 6 μg/mouse-treated mice, which were xenografted with the indicated H460 sgβ_2_AR-5/WT β_2_AR cells, lung tumors disappeared in 2 of 8 mice. The n-values denote the number of mice per group. SX: Salmeterol xinafoate. CR: Complete response. Scale bars: 2 mm. (**D**-**F**) Representative images of hematoxylin and eosin-stained lung tissue sections (D), change in the tumor number per lung between the control and salmeterol xinafoate treatment groups (E), and the fold change in tumor volume per lung between the control and salmeterol xinafoate treatment groups (F) on day 51 from BALB/c nude mice xenografted with the indicated H460 human lung cancer cells (1 × 10^6^) by intravenous tail vein injection. In endogenous β_2_AR knockout H460 lung cancer cells (sgβ_2_AR-6), wild-type human β_2_AR (WT β_2_AR), salmeterol xinafoate-binding-deficient human β_2_AR (F193W, F194V, S203T, S204T, H296K, R304P, K305D, and Y308F mutations, SBD β_2_AR), or G-protein-binding-deficient human β_2_AR (R131A and F139A mutations, GBD β_2_AR) was overexpressed as indicated. A once-weekly inhalational administration of salmeterol xinafoate (6 μg/mouse) was initiated from day 21 when the lung tumor was detected (Figure S1A). The experimental endpoint was on day 51 when the first dead mouse was observed in the control group. All lungs were analyzed by serial sectioning the entire lung and scoring the tumor nodules. A tumor nodule larger than 100 μm in width was counted and calculated as described in the Materials and Methods. Of the 6 μg/mouse-treated mice, which were xenografted with the indicated H460 sgβ_2_AR-6/WT β_2_AR cells, lung tumors disappeared in 2 of 8 mice. The n-values denote the number of mice per group. SX: Salmeterol xinafoate. CR: Complete response. Scale bars: 2 mm. (**G**) Change in the primary tumor volume between the control and salmeterol xinafoate treatment groups of BALB/c nude mice orthotopically xenografted with the indicated MDA-MB-231 human breast cancer cells (1 × 10^6^). In endogenous β_2_AR knockout MDA-MB-231 breast cancer cells (sgβ_2_AR-5), wild-type human β_2_AR (WT β_2_AR), salmeterol xinafoate-binding-deficient human β_2_AR (F193W, F194V, S203T, S204T, H296K, R304P, K305D, and Y308F mutations, SBD β_2_AR), or G-protein-binding-deficient human β_2_AR (R131A and F139A mutations, GBD β_2_AR) was overexpressed as indicated. A once-weekly intraperitoneal administration of salmeterol xinafoate (20 μg/kg) was initiated from day 15 when the primary tumor reached approximately 90 mm^3^. The green arrow indicates the time of intraperitoneal injection of salmeterol xinafoate. The primary tumor growth curves were stopped on day 45 when the first primary tumor reached 2000 mm^3^. The n-values denote the number of mice per group. SX: Salmeterol xinafoate. (**H**) Overall survival curves of BALB/c nude mice orthotopically xenografted with the indicated MDA-MB-231 human breast cancer cells (1 × 10^6^), which overexpressed the wild-type human β_2_AR (WT β_2_AR) in endogenous β_2_AR knockout MDA-MB-231 breast cancer cells (sgβ_2_AR-5). A once-weekly intraperitoneal administration of salmeterol xinafoate (20 μg/kg) was initiated from day 15 when the primary tumor reached approximately 90 mm^3^. Mice were considered to have expired when the primary tumor reached 2000 mm^3^. Of the 20 μg/kg-treated mice, tumors disappeared in 2 of 8 mice, and then lived for at least 139 and 124 days, without detectable tumors until the experimental endpoint at day 172. The colored number is the median survival time. The n-values denote the number of mice per group. SX: Salmeterol xinafoate. CR: Complete response. # The experimental endpoint on day 172. (**I**) Change in the primary tumor volume between the control and salmeterol xinafoate treatment groups of BALB/c nude mice orthotopically xenografted with the indicated MDA-MB-231 human breast cancer cells (1 × 10^6^). In endogenous β_2_AR knockout MDA-MB-231 breast cancer cells (sgβ_2_AR-6), wild-type human β_2_AR (WT β_2_AR), salmeterol xinafoate-binding-deficient human β_2_AR (F193W, F194V, S203T, S204T, H296K, R304P, K305D, and Y308F mutations, SBD β_2_AR), or G-protein-binding-deficient human β_2_AR (R131A and F139A mutations, GBD β_2_AR) was overexpressed as indicated. A once-weekly intraperitoneal administration of salmeterol xinafoate (20 μg/kg) was initiated from day 15 when the primary tumor reached approximately 90 mm^3^. The green arrow indicates the time of intraperitoneal injection of salmeterol xinafoate. The primary tumor growth curves were stopped on day 45 when the first primary tumor reached 2000 mm^3^. The n-values denote the number of mice per group. SX: Salmeterol xinafoate. (**J**) Overall survival curves of BALB/c nude mice orthotopically xenografted with the indicated MDA-MB-231 human breast cancer cells (1 × 10^6^), which overexpressed the wild-type human β_2_AR (WT β_2_AR) in endogenous β_2_AR knockout MDA-MB-231 breast cancer cells (sgβ_2_AR-6). A once-weekly intraperitoneal administration of salmeterol xinafoate (20 μg/kg) was initiated from day 15 when the primary tumor reached approximately 90 mm^3^. Mice were considered to have expired when the primary tumor reached 2000 mm^3^. Of the 20 μg/kg-treated mice, tumors disappeared in 3 of 9 mice, and then lived for at least 175, 148, and 139 days, without detectable tumors until the experimental endpoint at day 199. The colored number is the median survival time. The n-values denote the number of mice per group. SX: Salmeterol xinafoate. CR: Complete response. # The experimental endpoint on day 199. (**K**-**M**) Representative images of hematoxylin and eosin-stained lung tissue sections (K), change in the tumor number per lung between the control and salmeterol xinafoate treatment groups (L), and the fold change in tumor volume per lung between the control and salmeterol xinafoate treatment groups (M) on day 39 from C57BL/6 mice isografted with the indicated LLC mouse lung cancer cells (3 × 10^4^) by intravenous tail vein injection. In endogenous β_2_AR knockout LLC lung cancer cells (sgβ_2_AR-7), wild-type mouse β_2_AR (WT β_2_AR), salmeterol xinafoate-binding-deficient mouse β_2_AR (F193W, F194V, S203T, S204T, H296K, K305D, and Y308F mutations, SBD β_2_AR), or G-protein-binding-deficient mouse β_2_AR (R131A and F139A mutations, GBD β_2_AR) was overexpressed as indicated. A once-weekly inhalational administration of salmeterol xinafoate (6 μg/mouse) was initiated from day 5 when the lung tumor was detected (Figure S2A). The experimental endpoint was on day 39 when the first dead mouse was observed in the control group. All lungs were analyzed by serial sectioning the entire lung and scoring the tumor nodules. A tumor nodule larger than 100 μm in width was counted and calculated as described in the Materials and Methods. Of the 6 μg/mouse-treated mice, which were isografted with the indicated LLC sgβ_2_AR-7/WT β_2_AR cells, lung tumors disappeared in 2 of 8 mice. The n-values denote the number of mice per group. SX: Salmeterol xinafoate. CR: Complete response. Scale bars: 2 mm. (**N**-**P**) Representative images of hematoxylin and eosin-stained lung tissue sections (N), change in the tumor number per lung between the control and salmeterol xinafoate treatment groups (O), and the fold change in tumor volume per lung between the control and salmeterol xinafoate treatment groups (P) on day 39 from C57BL/6 mice isografted with the indicated LLC mouse lung cancer cells (3 × 10^4^) by intravenous tail vein injection. In endogenous β_2_AR knockout LLC lung cancer cells (sgβ_2_AR-8), wild-type mouse β_2_AR (WT β_2_AR), salmeterol xinafoate-binding-deficient mouse β_2_AR (F193W, F194V, S203T, S204T, H296K, K305D, and Y308F mutations, SBD β_2_AR), or G-protein-binding-deficient mouse β_2_AR (R131A and F139A mutations, GBD β_2_AR) was overexpressed as indicated. A once-weekly inhalational administration of salmeterol xinafoate (6 μg/mouse) was initiated from day 5 when the lung tumor was detected (Figure S2A). The experimental endpoint was on day 39 when the first dead mouse was observed in the control group. All lungs were analyzed by serial sectioning the entire lung and scoring the tumor nodules. A tumor nodule larger than 100 μm in width was counted and calculated as described in the Materials and Methods. Of the 6 μg/mouse-treated mice, which were isografted with the indicated LLC sgβ_2_AR-8/WT β_2_AR cells, lung tumors disappeared in 2 of 8 mice. The n-values denote the number of mice per group. SX: Salmeterol xinafoate. CR: Complete response. Scale bars: 2 mm. (**Q** and **R**) Change in the primary tumor volume between the control and salmeterol xinafoate treatment groups of BALB/c mice orthotopically isografted with the indicated 4T1 mouse breast cancer cells (1 × 10^6^). In endogenous β_2_AR knockout 4T1 breast cancer cells (sgβ_2_AR-7, Q or sgβ_2_AR-8, R), wild-type mouse β_2_AR (WT β_2_AR), salmeterol xinafoate-binding-deficient mouse β_2_AR (F193W, F194V, S203T, S204T, H296K, K305D, and Y308F mutations, SBD β_2_AR), or G-protein-binding-deficient mouse β_2_AR (R131A and F139A mutations, GBD β_2_AR) was overexpressed as indicated. A once-weekly intraperitoneal administration of salmeterol xinafoate (2 μg/kg) was initiated from day 7 (Q) or 8 (R) when the primary tumor reached approximately 100 mm^3^. The green arrow indicates the time of intraperitoneal injection of salmeterol xinafoate. The primary tumor growth curves were stopped on day 31 when the first dead mouse was observed in the control group. The n-values denote the number of mice per group. SX: Salmeterol xinafoate. Anesthetics were not used in the mouse experiments, except for cancer cells injection, because anesthetics prevented the anticancer effects of salmeterol xinafoate. Mice were fasted for 6 h (H460 and MDA-MB-231) or 12 h (LLC and 4T1) before salmeterol xinafoate administration. Four hours (H460 and MDA-MB-231) or 8 h (LLC and 4T1) after salmeterol xinafoate administration, food was provided. The data are presented as the mean ± s.e.m. values. *P*-values were determined by unpaired one-way ANOVA with uncorrected Fisher’s LSD test (B, C, E, F, L, M, O, and P), unpaired two-way ANOVA with uncorrected Fisher’s LSD test (G, I, Q, and R), or the log-rank test (H and J). \**P* < 0.05, \*\**P* < 0.01, \*\*\**P* < 0.001, and \*\*\*\**P* < 0.0001.

Furthermore, to establish whether β_2_AR downstream signaling is required for the anticancer effects of salmeterol xinafoate *in vivo*, both the mutant human β_2_AR with R131A and F139A mutations (GBD β_2_AR) and the mutant mouse β_2_AR with R131A and F139A mutations (GBD β_2_AR), which cannot induce downstream signaling (*79–81*), were used in tumorigenesis experiments in mice. In endogenous β_2_AR knockout cancer cells, the exogenous β_2_AR mutant with impaired downstream signaling completely prevented the anticancer effects of salmeterol xinafoate administered through inhalation (lung cancer) or intraperitoneal injection (breast cancer) in immunodeficient mice xenografted with human lung (Figures 3A-3F and S5) or breast cancer cells (Figures 3G-3J and S6) and in immunocompetent mice isografted with mouse lung (6 μg/mouse, Figures 3K-3P and S7) or breast cancer cells (2 μg/kg, Figures 3Q, 3R, and S8). Taken together, these findings demonstrate the requirement of the downstream signaling pathway for the anticancer effects of salmeterol xinafoate *in vivo*.

### Salmeterol xinafoate suppresses tumor sphere formation *in vitro*

Cancer stem cells drive the incidence, growth, and metastasis of cancers and have recently become novel drug targets for cancer therapies. We next conducted a tumorsphere formation assay to examine the effect of salmeterol xinafoate on the self-renewal of cancer stem cells *in vitro*. We found that treatment with salmeterol xinafoate dose-dependently decreased the capacity of lung cancer, breast cancer, and melanoma cells to form tumor spheres (Figures 4 and S9). These results indicate that salmeterol xinafoate exerts anticancer effects by inhibiting the self-renewal of cancer stem cells.

**Figure 4.**
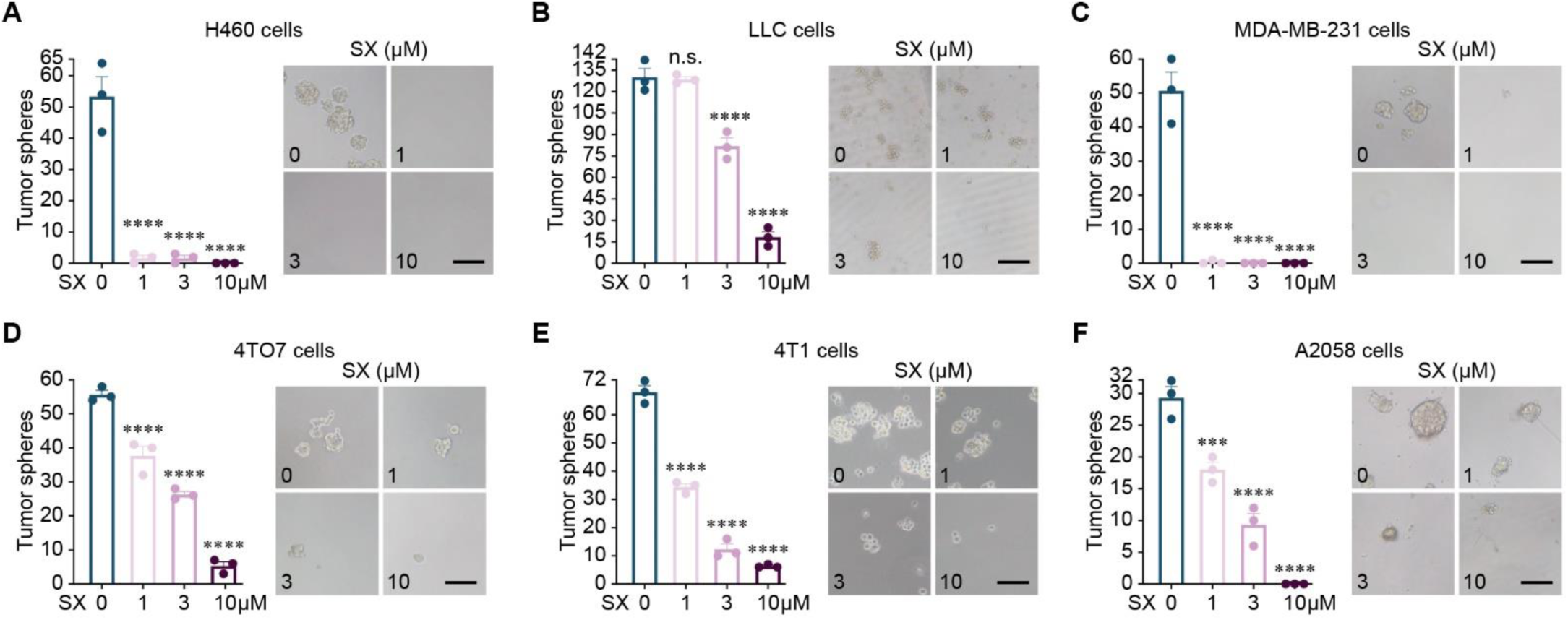
Salmeterol xinafoate suppresses the capacity of lung cancer, breast cancer, and melanoma cells to form tumor spheres. (**A** and **B**) Number of tumor spheres derived from H460 human lung cancer cells (400, A) and LLC mouse lung cancer cells (1,000, B) without or with salmeterol xinafoate treatment at the indicated doses (1, 3, or 10 μM) for 7 days (3 independent experiments, left). Representative images of tumor spheres derived from H460 human lung cancer cells (400, A) and LLC mouse lung cancer cells (1,000, B) without or with salmeterol xinafoate treatment at the indicated doses (1, 3, or 10 μM) for 7 days (right). SX: Salmeterol xinafoate. Scale bar: 100 μm. (**C**-**E**) Number of tumor spheres derived from MDA-MB-231 human breast cancer cells (1,000, C), 4TO7 mouse breast cancer cells (1,000, D), and 4T1 mouse breast cancer cells (1,000, E) without or with salmeterol xinafoate treatment at the indicated doses (1, 3, or 10 μM) for 7 days (3 independent experiments, left). Representative images of tumor spheres derived from MDA-MB-231 human breast cancer cells (1,000, C), 4TO7 mouse breast cancer cells (1,000, D), and 4T1 mouse breast cancer cells (1,000, E) without or with salmeterol xinafoate treatment at the indicated doses (1, 3, or 10 μM) for 7 days (right). SX: Salmeterol xinafoate. Scale bar: 100 μm. (**F**) Number of tumor spheres derived from A2058 human melanoma cells (1,000) without or with salmeterol xinafoate treatment at the indicated doses (1, 3, or 10 μM) for 7 days (3 independent experiments, left). Representative images of tumor spheres derived from A2058 human melanoma cells (1,000) without or with salmeterol xinafoate treatment at the indicated doses (1, 3, or 10 μM) for 7 days (right). SX: Salmeterol xinafoate. Scale bar: 100 μm. The data are presented as the mean ± s.e.m. values. *P*-values were determined by unpaired one-way ANOVA with uncorrected Fisher’s LSD test (A-F). \*\*\**P* < 0.001, \*\*\*\**P* < 0.0001, and n.s., not significant.

## Discussion

Several million years of biocultural evolution has shaped the bodies and minds of modern humans. In the evolutionary context, the hunter-gatherer way of life selected for the physical activity–sympathetic nervous system–catecholamines pathway to respond to a perceived threat from the external and internal environments of the body. In the context of cancer, accumulated evidence indicates that stimulation of β_2_AR (expressed on cancer cells by the above-described pathway) can control the initiation, growth, and metastasis of various cancers and improve the overall survival of patients (*56–59*). Because the evolutionarily selected physical activity–sympathetic nervous system–catecholamines–β_2_AR pathway has numerous beneficial effects on cancer, the identification of medicinal products that mimic or potentiate these effects is a longstanding medical goal (*3, 51–55*). In this study, we provide evidence that weekly administration of salmeterol xinafoate, a long-acting selective β_2_AR agonist drug, decreases cancer incidence, therapeutically inhibits cancer growth, and prolongs survival in mouse models of lung cancer, breast cancer, and melanoma. Our study reveals that in dealing with cancer, β_2_AR represents an ideal drug target and that salmeterol xinafoate monotherapy has strong anticancer activity. In addition, there is evidence to suggest that the physical activity–sympathetic nervous system– catecholamines–β_2_AR pathway enhances anticancer immunity by changing the magnitude and quality of immune system activity, such as by increasing the efflux of CD8^+^ T cells and NK cells into the circulating blood, regulating the function of CD8^+^ and γδ T cells, and diminishing the immunosuppressive effects of myeloid-derived suppressor cells on T cells (*28, 82–87*). Recently, immune checkpoint blockade therapies have shown extraordinary clinical effectiveness in patients with advanced cancer by increasing the anticancer activity of CD8^+^ T cells. Therefore, considering the strong effects on both cancer cells and immune cells, including CD8^+^ T cells, we speculated that the evolutionarily selected pathway plays a central role in cancer pathogenesis and therapy.

Inhaled medications are a unique treatment option and the optimal route of drug delivery for treating respiratory diseases because of their direct drug-pharmacological target interactions (*73–76*). However, currently, there are no inhaled drugs available for lung cancer therapy. In this study, we found that a once-weekly inhalation of salmeterol xinafoate exerted anticancer effects on lung cancer *in vivo* as effectively as a once-weekly intraperitoneal injection of salmeterol xinafoate. Intriguingly, even on breast cancer, similar anticancer effects were observed for a once-weekly inhalation of salmeterol xinafoate. Thus, we presume that inhaled medication represents a powerful and new route of drug delivery for the treatment of cancers, including but not limited to lung cancer.

G-protein-coupled receptors serve as the largest class of drug targets and 135 G-protein-coupled receptors are targeted by 700 drugs for clinical use, accounting for 35% of the marketed drugs (*88*). Recently, multiple lines of evidence have suggested the existence of multiple conformations of the active state of G-protein-coupled receptors that have various intrinsic efficacies for different signaling pathways. Hence, biased agonists are described as the preferential stabilization of unique conformations of agonist-activated G-protein-coupled receptors, which can selectively activate only one or a subgroup of downstream signaling pathways (*61, 62, 89–92*). Because activation of different signaling pathways can induce distinct biological functions, this biased agonism phenomenon is revolutionizing drug discovery for G-protein-coupled receptors and may also be used to identify novel therapeutic applications for existing approved drugs beyond the original medical indications (drug repurposing or repositioning) (*89, 91, 93*). Interestingly, previous studies have indicated that salmeterol xinafoate, a selective β_2_AR agonist drug, shows a 5- to 20-fold bias toward the activation of Gα_s_-mediated signaling over the stimulation of β-arrestin-mediated effects (*61–63, 70*). However, whether salmeterol xinafoate-induced biased signaling of β_2_AR is critical for its anticancer effects needs to be investigated further. In addition, as the first long-acting β_2_AR agonist used clinically and the most prescribed drug used as a bronchodilator for the treatment of asthma and chronic obstructive pulmonary disease, the pharmacokinetics and toxicity of salmeterol xinafoate are clearly understood (*67, 70, 73–75, 77, 78*). Moreover, the salmeterol xinafoate doses used in our mouse experiments are comparable to those used in patients with asthma or chronic obstructive pulmonary disease (Tables S4 and S5). Collectively, we propose that salmeterol xinafoate represents a promising therapeutic β_2_AR agonist and may be ready to enter clinical trials for patients with lung cancer, breast cancer, and melanoma.

## Author contributions

X.S. and J.M. designed, performed, and analyzed most experiments. H.G. conceived, designed, interpreted, and supervised the study, and wrote the manuscript with input from X.S. and J.M.

## Acknowledgments

We thank Drs. F. Giancotti (Columbia University) and P. Wang (Tongji University) for reagents, Drs. H. Qin (Shanghai Tenth People’s Hospital, Tongji University) and F. Sun (Shanghai Tenth People’s Hospital, Tongji University) for support and help, and members of the Gao laboratory for comments. This work was supported by the startup funding from Shanghai Tenth People’s Hospital to H.G.

