## Supplemental material for "Weekly inhaled salmeterol xinafoate, a selective β2 adrenergic receptor agonist drug, therapeutically inhibits cancer growth and prolongs survival *in vivo*"

#### Materials and methods

##### Cell lines

The H460 (K-Ras Q61H), H1299 (N-Ras Q61K/p53del), and H446 (p53 G154V/c-myc amplified) human lung cancer cell lines, the MDA-MB-231 (triple-negative/K-Ras G13D/p53 R280K) human breast cancer cell line, and the A2058 (p53 V274F/BRAF V600E) and A375 (BRAF V600E) human melanoma cell lines were obtained from the American Type Culture Collection (ATCC).

The LLC cell line (originating from a Lewis lung carcinoma in a C57BL/6 mouse) was originally obtained from the ATCC. LLC cells harbor a heterozygous K-Ras G12V mutation (*1*). The 4T1 cell line (triple-negative/CDKN2Adel/Kit A942S/p53 P31X, originating from a mammary carcinoma in a BALB/c mouse) (*2*) was obtained from the ATCC. The 4TO7 cell line (triple-negative, originating from a mammary carcinoma in a BALB/c mouse) was provided by Dr. Filippo G. Giancotti (Columbia University). The B16F10 cell line (Ink4a/Arf exons 1α 1β and 2 del, originating from melanoma in a C57BL/6 mouse) (*3*) was obtained from the ATCC. The Yummer 1.7 cell line (BRAF V600E/PTENdel/CDKN2Adel, originating from melanoma in a C57BL/6 mouse) was provided by Dr. Ping Wang (Tongji University).

The 293FT cell line was purchased from Life Technologies.

H460, H1299, and H446 cells were cultured in RPMI 1640 medium supplemented with 10% fetal bovine serum (FBS, A0500-3010, Cegrogen), 2 mM L-glutamine (21051-024, Gibco), and 100 U/ml penicillin / 0.1 mg/ml streptomycin (C0222, Beyotime Biotechnology). MDA-MB-231, A2058, A375, 293FT, LLC, 4T1, 4TO7, B16F10, and Yummer 1.7 cells were cultured in DMEM-HG supplemented with 10% FBS, 2 mM L-glutamine, and 100 U/ml penicillin / 0.1 mg/ml streptomycin.

All human cell lines were authenticated by short tandem repeat (STR) analysis provided by the Bio-Research Innovation Center Suzhou, SIBCB, CAS, and were routinely tested for mycoplasma contamination.

**Mice**

Mice were housed under specific pathogen-free (SPF) conditions in the animal facility of Tongji University. All animal experiments were approved by the Institutional Animal Care and Use Committee of Tongji University (approval number: TJBA01022101). BALB/c nude, C57BL/6, and BALB/c mice were purchased from the Shanghai SLAC Laboratory Animal Center (Shanghai, China). Mouse mammary tumor virus-polyoma middle T antigen (MMTV-PyMT) mice (on an FVB background) were purchased from Gempharmatech and bred at Tongji University.

H460 and A2058 cells were xenografted into 5- to 7-week-old male BALB/c nude mice. MDA-MB-231 cells were xenografted into 5- to 7-week-old female BALB/c nude mice. LLC and Yummer 1.7 cells were isografted into 5- to 7-week-old male syngeneic C57BL/6 mice. 4T1 and 4TO7 cells were isografted into 5- to 7-week-old female syngeneic BALB/c mice.

Anesthetics (such as isoflurane, ketamine-xylazine, and pentobarbital sodium, etc.) were not used in the mouse experiments, except for cancer cells injection, because anesthetics prevented the anticancer effects of salmeterol xinafoate.

**Lung cancer models**

Anesthetics (such as isoflurane, ketamine-xylazine, and pentobarbital sodium, etc.) were not used in the mouse experiments, except for cancer cells injection, because anesthetics prevented the anticancer effects of salmeterol xinafoate.

Cancer cells were harvested by trypsinization, washed once in phosphate-buffered saline (PBS), resuspended at 1 × $10^6$ (H460 or H460-Cas9 cells transduced with sgβ$_2$AR-5/WT β$_2$AR, sgβ$_2$AR-5/SBD β$_2$AR, sgβ$_2$AR-5/GBD β$_2$AR, sgβ$_2$AR-6/WT β$_2$AR, sgβ$_2$AR-6/SBD β$_2$AR, or sgβ$_2$AR-6/GBD β$_2$AR), 3 × $10^4$ (LLC), 3 × $10^4$ (LLC-Cas9 cells transduced with sgβ$_2$AR-7/WT β$_2$AR, sgβ$_2$AR-7/SBD β$_2$AR, sgβ$_2$AR-7/GBD β$_2$AR, sgβ$_2$AR-8/WT β$_2$AR, sgβ$_2$AR-8/SBD β$_2$AR, or sgβ$_2$AR-8/GBD β$_2$AR) cells in 50 μl of PBS and injected into mice by intravenous tail vein injection on day 0. A once-weekly administration of salmeterol xinafoate (S4296, Selleck) was initiated from day 21 (H460) or day 5 (LLC) when the lung tumor was detected. Mice were fasted for 6 h (H460) or 12 h (LLC) before salmeterol xinafoate administration. Four hours (H460) or 8 h (LLC) after salmeterol xinafoate administration, food was provided. Mice were randomized before grouping. Mice were sacrificed when they developed emaciation, lethargy, or irritability.

**Breast orthotopic cancer models**

Anesthetics (such as isoflurane, ketamine-xylazine, and pentobarbital sodium, etc.) were not used in the mouse experiments, except for cancer cells injection, because anesthetics prevented the anticancer effects of salmeterol xinafoate.

Cancer cells were harvested by trypsinization, washed once in PBS, resuspended at 1 × $10^6$ (MDA-MB-231 or MDA-MB-231-Cas9 cells transduced with sgβ$_2$AR-5/WT β$_2$AR, sgβ$_2$AR-5/SBD β$_2$AR, sgβ$_2$AR-5/GBD β$_2$AR, sgβ$_2$AR-6/WT β$_2$AR, sgβ$_2$AR-6/SBD β$_2$AR, or sgβ$_2$AR-6/GBD β$_2$AR) cells in 30 μl of 1:1 DMEM-HG/growth factor-reduced Matrigel (354234, Corning), and injected into the 4$^{th}$ mammary fat pad on the left side of mice on day 0. A once-weekly administration of salmeterol xinafoate was initiated from day 15, when the primary tumor reached approximately 90 $mm^3$. Mice were fasted for 6 h before salmeterol xinafoate administration. Four hours after salmeterol xinafoate administration, food was provided. The primary tumor growth was monitored by measuring the tumor length (L) and width (W). The tumor volume was calculated using the formula $LW^2/2$. Mice were sacrificed when the primary tumor reached 2000 $mm^3$.

Cancer cells were harvested by trypsinization, washed once in PBS, resuspended at $1 \times 10^6$ (4TO7, 4T1, or 4T1-Cas9 cells transduced with sg$\beta_2$AR-7/WT $\beta_2$AR, sg$\beta_2$AR-7/SBD $\beta_2$AR, sg$\beta_2$AR-7/GBD $\beta_2$AR, sg$\beta_2$AR-8/WT $\beta_2$AR, sg$\beta_2$AR-8/SBD $\beta_2$AR, or sg$\beta_2$AR-8/GBD $\beta_2$AR) cells in 30 μl of DMEM-HG and injected into the 4$^{th}$ mammary fat pad on the left side of mice on day 0. A once-weekly administration of salmeterol xinafoate was initiated from day 7 or 8, when the primary tumor reached approximately 50 (4TO7) or 100 (4T1) $mm^3$. Mice were fasted for 12 h before salmeterol xinafoate administration. Eight hours after salmeterol xinafoate administration, food was provided. The primary tumor growth was monitored by measuring the tumor length (L) and width (W). The tumor volume was calculated using the formula $LW^2/2$. Mice were sacrificed when they developed emaciation, lethargy, or irritability.

**Melanoma orthotopic cancer models**

Anesthetics (such as isoflurane, ketamine-xylazine, and pentobarbital sodium, etc.) were not used in the mouse experiments, except for cancer cells injection, because anesthetics prevented the anticancer effects of salmeterol xinafoate.

Cancer cells were harvested by trypsinization, washed once in PBS, resuspended at $1 \times 10^6$ (A2058) and $3 \times 10^5$ (Yummer 1.7) cells in 30 μl of DMEM-HG, and injected subcutaneously into mice on day 0. A once-weekly administration of salmeterol xinafoate was initiated from day 11 or 14 when the primary tumor reached approximately 50 or 100 $mm^3$ (for A2058), or initiated from day 6 or 10 when the primary tumor reached approximately 50 $mm^3$ (for Yummer 1.7). Mice were fasted for 6 h (A2058) or 12 h (Yummer 1.7) before salmeterol xinafoate administration. Four hours (A2058) or 8 h (Yummer 1.7) after salmeterol xinafoate administration, food was provided. The primary tumor growth was monitored by measuring the tumor length (L) and width (W). The tumor volume was calculated using the formula $LW^2/2$. Mice were sacrificed when the primary tumor reached 2000 $mm^3$.

**MMTV-PyMT spontaneous breast cancer model**

Anesthetics (such as isoflurane, ketamine-xylazine, and pentobarbital sodium, etc.) were not used in the mouse experiments because anesthetics prevented the anticancer effects of salmeterol xinafoate.

Only virgin female MMTV-PyMT mice (FVB strain) were used. Before the tumor became palpable (28 days of age), when the tumor established (49 days of age), or when lung metastasis established (70 days of age), MMTV-PyMT mice were randomly assigned to the control or salmeterol xinafoate-treated groups, and tumor growth was measured weekly thereafter. Mice were fasted for 12 h before salmeterol xinafoate administration. Eight hours after salmeterol xinafoate administration, food was provided. The primary tumor growth was monitored by measuring the tumor length (L) and width (W). The tumor volume was calculated using the formula $LW^2/2$. Because MMTV-PyMT mice develop multiple mammary tumors, the total tumor volume was calculated as the sum of the volumes of all tumors per mouse. Mice were sacrificed when the total tumor volume reached 2000 $mm^3$.

**Intraperitoneal administration of salmeterol xinafoate**

Anesthetics (such as isoflurane, ketamine-xylazine, and pentobarbital sodium, etc.) were not used in the mouse experiments, except for cancer cells injection, because anesthetics prevented the anticancer effects of salmeterol xinafoate.

For the lung cancer models, the lung tumors were confirmed by immunofluorescence staining on day 21 from when BALB/c nude mice were xenografted with H460 human lung cancer cells or on day 5 from when C57BL/6 mice were isografted with LLC mouse lung cancer cells through intravenous tail vein injection. A once-weekly intraperitoneal administration of salmeterol xinafoate (2 or 20 μg/kg) or vehicle control (0.135% DMSO in PBS) was initiated from day 21 (for H460) or day 5 (for LLC) until the day of euthanasia. Mice were fasted for 6 h (H460) or 12 h (LLC) before intraperitoneal administration. Four hours (H460) or 8 h (LLC) after intraperitoneal administration, food was provided. Mice were randomized before grouping.

For the breast orthotopic cancer models, a once-weekly intraperitoneal administration of salmeterol xinafoate (2 or 20 μg/kg) or vehicle control (0.135% DMSO in PBS) was initiated from day 15, when the primary tumor reached approximately 90 $mm^3$ (for MDA-MB-231), or from day 7, when the primary tumor reached approximately 50 $mm^3$ (for 4TO7) or 100 $mm^3$ (for 4T1). The once-weekly intraperitoneal administration was stopped on the day when the primary tumor reached 2000 $mm^3$ (for MDA-MB-231 and 4TO7) or the day of euthanasia (for 4T1). Mice were fasted for 6 h (MDA-MB-231) or 12 h (4T1 or 4TO7) before intraperitoneal administration. Four hours (MDA-MB-231) or 8 h (4T1 or 4TO7) after intraperitoneal administration, food was provided. Mice were randomized before grouping.

A once-weekly intraperitoneal administration of salmeterol xinafoate (20 μg/kg) or vehicle control (0.135% DMSO in PBS) was initiated from day 15 when the primary tumor reached approximately 90 $mm^3$ (MDA-MB-231, MDA-MB-231-Cas9 cells transduced with sg$\beta_2$AR-5/WT $\beta_2$AR, sg$\beta_2$AR-5/SBD $\beta_2$AR, sg$\beta_2$AR-5/GBD $\beta_2$AR, sg$\beta_2$AR-6/WT $\beta_2$AR, sg$\beta_2$AR-6/SBD $\beta_2$AR, or sg$\beta_2$AR-6/GBD $\beta_2$AR). The once-weekly intraperitoneal administration was stopped on the day when the primary tumor reached 2000 $mm^3$. Mice were fasted for 6 h before intraperitoneal administration. Four hours after intraperitoneal administration, food was provided. Mice were randomized before grouping.

A once-weekly intraperitoneal administration of salmeterol xinafoate (2 μg/kg) or vehicle control (0.135% DMSO in PBS) was initiated from day 7 or 8 when the primary tumor reached approximately 100 $mm^3$ (4T1-Cas9 cells transduced with sgβ2AR-7/WT β2AR, sgβ2AR-7/SBD β2AR, sgβ2AR-7/GBD β2AR, sgβ2AR-8/WT β2AR, sgβ2AR-8/SBD β2AR, or sgβ2AR-8/GBD β2AR). The once-weekly intraperitoneal administration was stopped on the day of euthanasia. Mice were fasted for 12 h before intraperitoneal administration. Eight hours after intraperitoneal administration, food was provided. Mice were randomized before grouping.

For the melanoma orthotopic cancer models, a once-weekly intraperitoneal administration of salmeterol xinafoate (6 or 20 μg/kg for A2058 cells and 2 or 20 μg/kg for Yummer 1.7 cells) or vehicle control (0.135% DMSO in PBS) was initiated from day 11, when the primary tumor reached approximately 50 $mm^3$ (for A2058), or day 10, when the primary tumor reached approximately 50 $mm^3$ (for Yummer 1.7). The once-weekly intraperitoneal administration was stopped on the day when the primary tumor reached 2000 $mm^3$. Mice were fasted for 6 h (A2058) or 12 h (Yummer 1.7) before intraperitoneal administration. Four hours (A2058) or 8 h (Yummer 1.7) after intraperitoneal administration, food was provided. Mice were randomized before grouping.

For MMTV-PyMT spontaneous breast cancer model, a once-weekly intraperitoneal administration of salmeterol xinafoate (2 or 20 μg/kg) or vehicle control (0.135% DMSO in PBS) was initiated before the tumor became palpable (28 days of age), when the tumor became established (49 days of age), or when lung metastasis occurred (70 days of age). The once-weekly intraperitoneal administration was stopped on the day when the total mammary tumor reached 2000 $mm^3$. Mice were fasted for 12 h before intraperitoneal administration. Eight hours after intraperitoneal administration, food was provided. Mice were randomized before grouping. Mice were randomized before grouping.

**Inhalation administration of salmeterol xinafoate**

Anesthetics (such as isoflurane, ketamine-xylazine, and pentobarbital sodium, etc.) were not used in the mouse experiments, except for cancer cells injection, because anesthetics prevented the anticancer effects of salmeterol xinafoate.

For the lung cancer models, the lung tumors were detected by immunofluorescence staining on day 21, when BALB/c nude mice were xenografted with H460 human lung cancer cells, or on day 5 from when C57BL/6 mice were isografted with LLC mouse lung cancer cells through intravenous tail vein injection. A once-weekly inhalation administration of salmeterol xinafoate (2 or 6 μg/mouse for H460 cells and 2 or 20 μg/mouse for LLC cells) or vehicle control (1% DMSO in normal saline) was initiated from day 21 (H460) or day 5 (LLC) to the day of euthanasia. Mice were fasted for 6 h (H460) or 12 h (LLC) before inhalational administration. Four hours (H460) or 8 h (LLC) after inhalational administration, food was provided. Mice were randomized before grouping. The airflow rate was 135 μl/min, and the mice inhaled for 30 s.

A once-weekly inhalation administration of salmeterol xinafoate (6 μg/mouse) or vehicle control (1% DMSO in normal saline) was intiated from day 21 (H460-Cas9 cells transduced with sg$\beta_2$AR-5/WT $\beta_2$AR, sg$\beta_2$AR-5/SBD $\beta_2$AR, sg$\beta_2$AR-5/GBD $\beta_2$AR, sg$\beta_2$AR-6/WT $\beta_2$AR, sg$\beta_2$AR-6/SBD $\beta_2$AR, or sg$\beta_2$AR-6/GBD $\beta_2$AR). The once-weekly inhalation administration was stopped on the day of euthanasia. Mice were fasted for 6 h before inhalational administration. Four hours after inhalational administration, food was provided. Mice were randomized before grouping. The airflow rate was 135 μl/min, and the mice inhaled for 30 s.

A once-weekly inhalation administration of salmeterol xinafoate (6 μg/mouse) or vehicle control (1% DMSO in normal saline) was initiated from day 5 (LLC-Cas9 cells transduced with sg$\beta_2$AR-7/WT $\beta_2$AR, sg$\beta_2$AR-7/SBD $\beta_2$AR, sg$\beta_2$AR-7/GBD $\beta_2$AR, sg$\beta_2$AR-8/WT $\beta_2$AR, sg$\beta_2$AR-8/SBD $\beta_2$AR, or sg$\beta_2$AR-8/GBD $\beta_2$AR). The once-weekly inhalation administration was stopped on the day of euthanasia. Mice were fasted for 12 h before inhalational administration. Eight hours after inhalational administration, food was provided. Mice were randomized before grouping. The airflow rate was 135 μl/min, and the mice inhaled for 30 s.

For the breast orthotopic cancer models, a once-weekly inhalation administration of salmeterol xinafoate (2 or 6 μg/mouse) or vehicle control (1% DMSO in normal saline) was initiated from day 15, when the primary tumor reached approximately 90 $mm^3$ (for MDA-MB-231), or from day 7, when the primary tumor reached approximately 50 $mm^3$ (for 4TO7) or 100 $mm^3$ (for 4T1). The once-weekly inhalation administration was stopped on the day when the primary tumor reached 2000 $mm^3$ (for MDA-MB-231 and 4TO7) or the day of euthanasia (for 4T1). Mice were fasted for 6 h (MDA-MB-231) or 12 h (4T1 or 4TO7) before inhalational administration. Four hours (MDA-MB-231) or 8 h (4T1 or 4TO7) after inhalational administration, food was provided. Mice were randomized before grouping. The airflow rate was 135 μl/min, and the mice inhaled for 30 s.

For the MMTV-PyMT spontaneous breast cancer model, a once-weekly inhalation administration of salmeterol xinafoate (2 or 6 μg/mouse) or vehicle control (1% DMSO in normal saline) was initiated before the tumor became palpable (28 days of age), when the tumor became established (49 days of age), or when lung metastasis occurred (70 days of age). The once-weekly inhalation administration was stopped on the day when the total mammary tumor reached 2000 $mm^3$. Mice were fasted for 12 h before inhalational administration. Eight hours after inhalational administration, food was provided. Mice were randomized before grouping. The airflow rate was 135 μl/min, and the mice inhaled for 30 s.

**Transdermal administration of salmeterol xinafoate**

Anesthetics (such as isoflurane, ketamine-xylazine, and pentobarbital sodium, etc.) were not used in the mouse experiments, except for cancer cells injection, because anesthetics prevented the anticancer effects of salmeterol xinafoate.

The ointment consisted of 8% glycerol, 20% Vaseline, 4% propylene glycol, and 68% $ddH_2O$. Salmeterol xinafoate was added to the ointment to be applied to the surface of the melanoma.

A once-weekly transdermal administration of salmeterol xinafoate (20 or 60 μg/mouse) or vehicle control (8% glycerol, 20% Vaseline, and 4% propylene glycol in $ddH_2O$) was initiated from day 14, when the primary tumor reached approximately 100 $mm^3$ (for A2058), or day 6, when the primary tumor reached approximately 50 $mm^3$ (for Yummer 1.7). Mice were fasted for 6 h (A2058) or 12 h (Yummer 1.7) before transdermal administration. Four hours (A2058) or 8 h (Yummer 1.7) after transdermal administration, food was provided. The once-weekly transdermal administration was stopped on the day when the primary tumor reached 2000 $mm^3$. Mice were randomized before grouping.

**Tumor sphere formation assay**

Cancer cells were seeded in 96-well ultralow attachment plates. All tumor spheres in each well were counted. Three independent experiments were conducted.

The H460 (400/well), H1299 (1,000/well) human lung cancer cells, and LLC (1,000/well) mouse lung cancer cells were cultured for 7 days in DMEM/F12 (12500-062, Gibco) supplemented with 1 × B27 (17504-044, Invitrogen), 20 ng/ml epidermal growth factor (EGF, 11140050, Invitrogen), 10 ng/ml basic fibroblast growth factor (bFGF, PHG0261, Invitrogen), 2 μg/ml heparin (H3149-500KU-9, Sigma), 1 × non-essential amino acid (NEAA, 11140050, Invitrogen), and 100 U/ml penicillin / 0.1 mg/ml streptomycin (*4*) with or without salmeterol xinafoate (1, 3, 10 μM).

The H446 (3,000/well) human lung cancer cells were cultured for 7 days in DMEM/F12 supplemented with 1 × B27, 20 ng/ml EGF, 20 ng/ml bFGF, 5 μg/ml Insulin (I9278, Sigma), 0.4% BSA (B2064, Sigma), 1 × NEAA, and 100 U/ml penicillin / 0.1 mg/ml streptomycin (*5*) with or without salmeterol xinafoate (1, 3, or 10 μM).

The MDA-MB-231 (1,000/well) human breast cancer cells were cultured for 7 days in MEBM (CC-3151, Lonza) supplemented with 1 × B27, 20 ng/ml EGF, 20 ng/ml bFGF, 4 μg/ml heparin, 5 μg/ml insulin, 0.5 μg/ml hydrocortisone (H0888, Sigma), and 100 U/ml penicillin / 0.1 mg/ml streptomycin (*6*) with or without salmeterol xinafoate (1, 3, or 10 μM).

The 4T1 (1,000/well) mouse breast cancer cells were cultured for 7 days in DMEM/F12 supplemented with 1 × B27, 20 ng/ml EGF, 20 ng/ml bFGF, 10 μg/ml insulin, 20 μg/ml hydrocortisone, and 100 U/ml penicillin / 0.1 mg/ml streptomycin (*7*) with or without salmeterol xinafoate (1, 3, or 10 μM).

The 4TO7 (1,000/well) mouse breast cancer cells were cultured for 7 days in DMEM/F12 supplemented with 1 × B27, 10 ng/ml EGF, 10 ng/ml bFGF, 4 μg/ml heparin, 5 μg/ml insulin, 20 μg/ml hydrocortisone, and 100 U/ml penicillin / 0.1 mg/ml streptomycin (*6*) with or without salmeterol xinafoate (1, 3, or 10 μM).

The A2058 (1,000/well), A375 (1,000/well) human melanoma cells, and B16F10 (1,000/well) mouse melanoma cells were cultured for 7 days in DMEM/F12 supplemented with 1 × B27, 20 ng/ml EGF, 10 ng/ml bFGF, and 100 U/ml penicillin / 0.1 mg/ml streptomycin (*7*) with or without salmeterol xinafoate (1, 3, or 10 μM).

The Yummer 1.7 (3,000/well) mouse melanoma cells were cultured for 7 days in DMEM/F12 supplemented with 1 × B27, 20 ng/ml EGF, 20 ng/ml bFGF, 4 μg/ml insulin, and 100 U/ml penicillin / 0.1 mg/ml streptomycin with or without salmeterol xinafoate (1, 3, or 10 μM).

**Hematoxylin and Eosin (H&E) staining and lung tumor statistics**

Mice bearing H460, H460-Cas9 cells transduced with sg$\beta_2$AR-5/WT $\beta_2$AR, sg$\beta_2$AR-5/SBD $\beta_2$AR, sg$\beta_2$AR-5/GBD $\beta_2$AR, sg$\beta_2$AR-6/WT $\beta_2$AR, sg$\beta_2$AR-6/SBD $\beta_2$AR, or sg$\beta_2$AR-6/GBD $\beta_2$AR human lung carcinomas were sacrificed on day 51 from when BALB/c nude mice were xenografted with H460 human lung cancer cells through intravenous tail vein injection (the day when the first dead mouse was observed in the control group).

Mice bearing LLC, LLC-Cas9 cells transduced with sg$\beta_2$AR-7/WT $\beta_2$AR, sg$\beta_2$AR-7/SBD $\beta_2$AR, sg$\beta_2$AR-7/GBD $\beta_2$AR, sg$\beta_2$AR-8/WT $\beta_2$AR, sg$\beta_2$AR-8/SBD $\beta_2$AR, or sg$\beta_2$AR-8/GBD $\beta_2$AR mouse lung carcinomas were sacrificed on day 39 from when

C57BL/6 mice were isografted with LLC mouse lung cancer cells through intravenous tail vein injection (the day when the first dead mouse was observed in the control group).

Lungs were fixed in 4% PFA for 36 h at 4°C, dehydrated in different concentrations of ethanol, followed by Histo-Clear® II (HS-202, National Diagnostics), and finally embedded in paraffin. The entire lung was serially sectioned (6-10 μm thick) using a Leica microtome. The sections were stained with hematoxylin (GS-02, Green Specimen) and eosin (G1001, Servicebio). The number of tumors was counted at 4 × magnification. A tumor nodule larger than 100 μm in width was counted. The cross-sections of each tumor on different sections were compared, and the largest cross-section of each tumor was chosen to measure the length and width of each tumor. Considering the paraffin-embedded shrinkage rate (40%), the length and width of the tumor were divided by 0.6, and the corrected length (L) and width (W) of the tumor were obtained. The tumor volume was calculated using the following formula: $LW^2/2$. Because mice develop multiple lung tumors, the total tumor volume was calculated as the sum of the volumes of all tumors per mouse. Representative images were scanned using a 3D Histech MIDI panoramic scanner.

**Immunofluorescence staining**

The mice were sacrificed on day 21 from when BALB/c nude mice were xenografted with H460 human lung cancer cells through intravenous tail vein injection (two mice) and on day 5 from C57BL/6 mice were isografted with LLC mouse lung cancer cells through intravenous tail vein injection (two mice).

The mice were then perfused with PBS through the left ventricle. Lungs were fixed in 4% PFA overnight at 4°C, dehydrated in 15% sucrose for 24 h, dehydrated in 30% sucrose for 24 h, and embedded in OCT compound. Nonconsecutive sections (10 μm thick) were sliced with a Leica microtome, and immunofluorescence staining was performed with goat anti-tdTomato antibody (orb182397, Biorbyt, RRID: AB_2687917) using a Tyramide Signal Amplification Kit (B40931, Thermo Fisher Scientific) to visualize the tumor cells and rabbit anti-CD31 antibody (Ab28364, Abcam, RRID: AB_726362) to visualize the lung capillaries. Immunoreactions were detected using fluorescently labeled secondary antibodies (Alexa Fluor$^{TM}$ 488-anti-rabbit IgG, A11034, Thermo Fisher Scientific, RRID: AB_2576217; Alexa Fluor$^{TM}$ 568-Streptavidin, S11226, Thermo Fisher Scientific, RRID: AB_144696; and Biotin-anti-goat IgG, BA-9500, Vector Labs, RRID: AB_2336123). Sections were mounted using ProLong$^{TM}$ Gold Antifade Mountant with DAPI (P36931, Thermo Fisher Scientific) and imaged using a Leica fluorescence microscope.

**Analysis of protein expression**

The Nucl-Cyto-Mem Preparation Kit (P1201, Applygen) was used to prepare the cell membrane, which was lysed in 1% Triton X-100/TBS (10 mM Tris-HCl pH 7.4, 0.9% NaCl, 0.02% KCl) supplemented with $Na_3VO_4$, phosphatase inhibitor cocktail (B15001, Bimake), and protease inhibitors (539134, Merck). The Enhanced BCA Protein Assay Kit was used to measure the protein concentration. Protein expression was determined by immunoblotting with primary antibodies (mouse anti-human $\beta_2$AR, clone: E-3, sc-271322, Santa Cruz, RRID: AB_10610813; mouse anti-mouse $\beta_2$AR, clone: R11E1, sc-81577, Santa Cruz, RRID: AB_1119478; mouse anti-HA, clone: 2S8Z1, AE008, ABclonal, RRID: AB_2770404; and rabbit anti-ATP1A1, 55187-1-AP, Proteintech, RRID: AB_10859261) and secondary antibodies (HRP-anti-Mouse IgG, BF03001, Blodragon; and HRP-anti-Rabbit IgG, BF03008, Blodragon).

**Liquid chromatograph-mass spectrometry (LC-MS)**

In the MMTV-PyMT mouse model of spontaneous breast cancer, 7-week-old female mice with established tumors were administered a once-weekly intraperitoneal injection of salmeterol xinafoate (20 μg/kg). Mice were fasted for 12 h before inhalation administration. The mice were sacrificed at 10, 30, 60, 120, and 240 min after the third injection, and breast tumors (1-3 tumors) were collected. The tumor weight was calculated as the sum of the weight of all tumors per mouse. All tumors from one mouse were mechanically minced together and taken as one sample for LC-MS analysis (Nanjing Biorn Lifescience Co., Ltd).

**Plasmids**

Constructs encoding sgRNAs against human $\beta_2$AR (#5, GCAACTTCTGGTGCGAGTTT and #6, ACCACGACGTCACGCAGGAA) and mouse $\beta_2$AR (#7, ACGGGACGAAGCGTGGGTTG and #8, TCTGGCGCTCGGCTTCCGTT) were generated by cloning the corresponding sgRNA sequences into the lentiGuide-Puro vector, which was a gift from Feng Zhang (Addgene plasmid #52963; http://n2t.net/addgene:52963; RRID: Addgene_52963).

cDNAs encoding full-length mouse and human $\beta_2$AR were cloned and sequenced. The human salmeterol xinafoate-binding-deficient mutant $\beta_2$AR (SBD) had site mutations including F193W, F194V, S203T, S204T, H296K, R304P, K305D, and Y308F. The human G-protein-binding-deficient mutant $\beta_2$AR (GBD $\beta_2$AR) had site mutations consisting of R131A and F131A (*8*). The mouse salmeterol xinafoate-binding-deficient mutant $\beta_2$AR (SBD) had site mutations including F193W, F194V, S203T, S204T, H296K, K305D, and Y308F. The mouse G-protein-binding-deficient mutant $\beta_2$AR (GBD $\beta_2$AR) had site mutations consisting of R131A and F131A (*8*).

Both human and mouse HA-WT β$_2$AR, HA-SBD β$_2$AR, and HA-GBD β$_2$AR were all resistant to sgRNAs for human or mouse β$_2$AR, and were subcloned into pQCXIN (Clontech) retroviral vectors and verified by sequencing.

Populations of knockout or overexpressing cells were used, and the efficiencies of knockout or overexpression were verified by western blotting.

**Statistical analysis**

Graphing and data analyses were performed using GraphPad Prism (version 8.3.0). The mean ± s.e.m. values are presented unless stated otherwise. The statistical methods and the numbers of replicates are listed in the figure legends. *P*-values <0.05 were considered statistically significant for all analyses.

Because 0 values could not be used to calculate log values, 0 was replaced with 1 when calculating the number of lung tumors in Supplementary Figures S5C (H460 sgβ$_2$AR-5/WT β$_2$AR SX 6 μg/mouse group) and S5F (H460 sgβ$_2$AR-6/WT β$_2$AR SX 6 μg/mouse group).

Because values of 0 could not be used to calculate log values and could not be used as a divisor, 0 mm$^3$ was replaced with 0.01 mm$^3$, which was approximately equal to the smallest lung tumor volume except for 0 mm$^3$ in Figures 3C and S5D (H460 sgβ$_2$AR-5/WT β$_2$AR SX 6 μg/mouse group).

Because values of 0 could not be used to calculate log values and could not be used as a divisor, 0 mm$^3$ was replaced with 0.001 mm$^3$, which was approximately equal to the smallest lung tumor volume except for 0 mm$^3$ in Figures 3F and S5G (H460 sgβ$_2$AR-6/WT β$_2$AR SX 6 μg/mouse group).

Because values of 0 could not be used as a divisor, 0 $mm^3$ was replaced with 0.151 $mm^3$, which was approximately equal to the smallest lung tumor volume except for 0 $mm^3$ in Figure 3M (LLC sg$\beta_2$AR-7/WT $\beta_2$AR SX 6 μg/mouse group).

Because values of 0 could not be used as a divisor, 0 $mm^3$ was replaced with 0.002 $mm^3$, which was approximately equal to the smallest lung tumor volume except for 0 $mm^3$ in Figure 3P (LLC sg$\beta_2$AR-8/WT $\beta_2$AR SX 6 μg/mouse group).

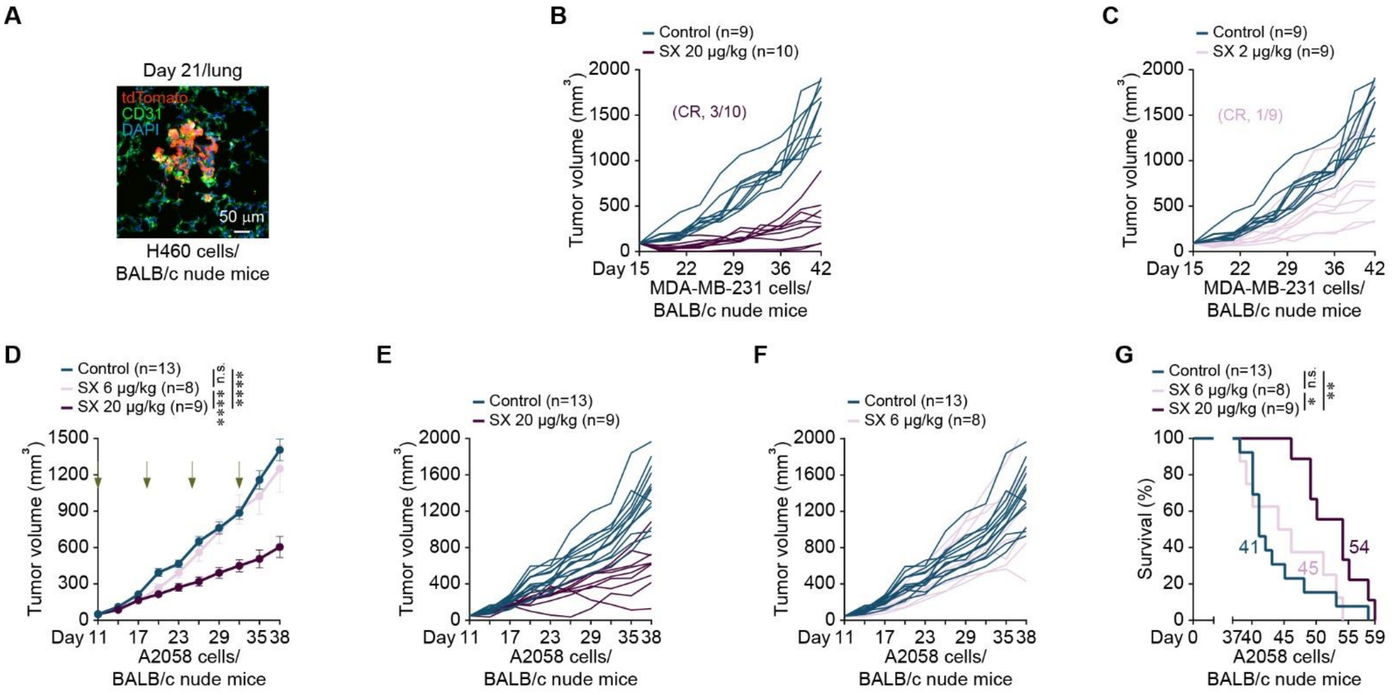


**Figure S1**

**Figure S1. A once-weekly intraperitoneal administration of salmeterol xinafoate therapeutically inhibits the growth of established tumors and prolongs survival of model mice with human lung cancer, breast cancer, and melanoma**

(**A**) Representative immunofluorescence image of a lung tissue section at day 21 from when BALB/c nude mice were xenografted with H460 human lung cancer cells ($1 \times 10^6$) by intravenous tail vein injection. tdTomato: H460 cells, CD31: Endothelial cells, DAPI: Cell nuclei.

(**B** and **C**) Primary tumor growth curves of individual BALB/c nude mice orthotopically xenografted with MDA-MB-231 human breast cancer cells ($1 \times 10^6$). A once-weekly intraperitoneal administration of salmeterol xinafoate (20 μg/kg, B or 2 μg/kg, C) was initiated from day 15 when the primary tumor reached approximately 90 $mm^3$. The primary tumor growth curves were stopped on day 42 when the first primary tumor reached 2000 $mm^3$. Of the 20 μg/kg-treated mice, tumors disappeared in 3 of 10 mice, and the 3 mice lived for at least 30, 45, and 57 days, without detectable tumors until the experimental endpoint at day 90. Of the 2 μg/kg-treated mice, tumors disappeared in 1 of 9 mice, and the mice lived for at least 33 days without detectable tumors until the experimental endpoint at day 90. The n-values denote the number of mice per group. SX: Salmeterol xinafoate. CR: Complete response.

(**D** and **G**) Primary tumor growth curves (D) and overall survival curves (G) of BALB/c nude mice orthotopically xenografted with A2058 human melanoma cells ($1 \times 10^6$). A once-weekly intraperitoneal administration of salmeterol xinafoate (6 or 20 μg/kg) was initiated from day 11 when the primary tumor reached approximately 50 $mm^3$. The green arrow indicates the time of intraperitoneal injection of salmeterol xinafoate. The primary tumor growth curves were stopped on day 38 when the first primary tumor reached 2000 $mm^3$ in the control group (D). Mice were considered to have expired when the primary tumor reached 2000 $mm^3$ (G). The colored number is the median survival time (G). The n-values denote the number of mice per group. SX: Salmeterol xinafoate.

(**E** and **F**) Primary tumor growth curves of individual BALB/c nude mice orthotopically xenografted with A2058 human melanoma cells ($1 \times 10^6$). A once-weekly intraperitoneal administration of salmeterol xinafoate (20 μg/kg, E or 6 μg/kg, F) was initiated from day 11 when the primary tumor reached approximately 50 $mm^3$. The primary tumor growth curves were stopped on day 38 when the first primary tumor reached 2000 $mm^3$ in the control group. The n-values denote the number of mice per group. SX: Salmeterol xinafoate.

Anesthetics were not used in the mouse experiments, except for cancer cells injection, because anesthetics prevented the anticancer effects of salmeterol xinafoate. Mice were fasted for 6 h before salmeterol xinafoate administration. Four hours after salmeterol xinafoate administration, food was provided. The data are presented as the mean ± s.e.m. values. *P*-values were determined by unpaired two-way ANOVA with uncorrected Fisher's least significant difference (LSD) test (D) or the log-rank test (G). *$P < 0.05$, **$P < 0.01$, ****$P < 0.0001$, and n.s., not significant.

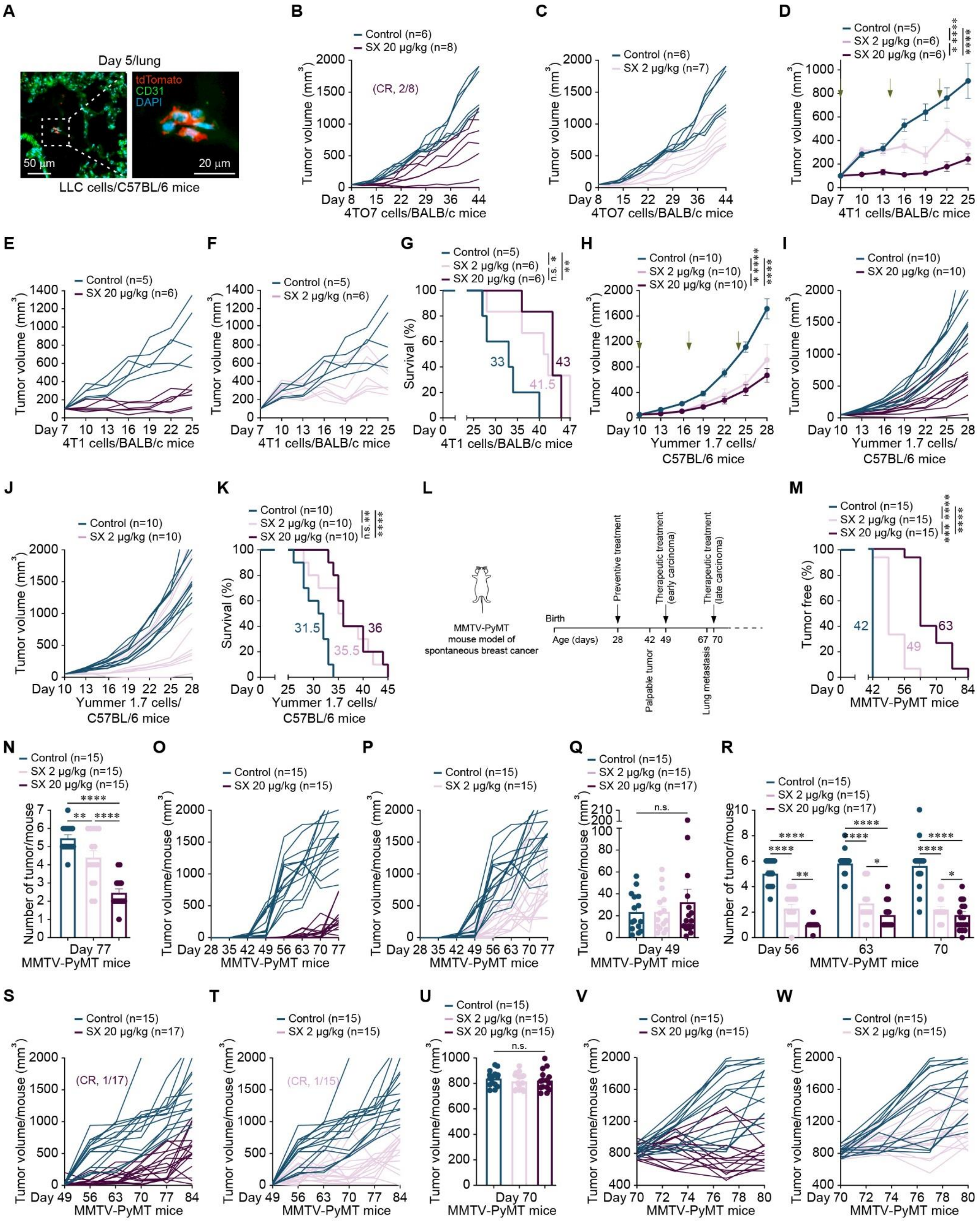
A
Day 5/lung
tdTomato
CD31
DAPI
50 µm
20 µm
LLC cells/C57BL/6 mice
B
Control (n=6)
SX 20 µg/kg (n=8)
(CR, 2/8)
Tumor volume (mm3)
4TO7 cells/BALB/c mice
C
Control (n=6)
SX 2 µg/kg (n=7)
4TO7 cells/BALB/c mice
D
Control (n=5)
SX 2 µg/kg (n=6)
SX 20 µg/kg (n=6)
4T1 cells/BALB/c mice
E
Control (n=5)
SX 20 µg/kg (n=6)
4T1 cells/BALB/c mice
F
Control (n=5)
SX 2 µg/kg (n=6)
4T1 cells/BALB/c mice
G
Control (n=5)
SX 2 µg/kg (n=6)
SX 20 µg/kg (n=6)
Survival (%)
33
41.5
43
4T1 cells/BALB/c mice
H
Control (n=10)
SX 2 µg/kg (n=10)
SX 20 µg/kg (n=10)
Yummer 1.7 cells/ C57BL/6 mice
I
Control (n=10)
SX 20 µg/kg (n=10)
Yummer 1.7 cells/ C57BL/6 mice
J
Control (n=10)
SX 2 µg/kg (n=10)
Yummer 1.7 cells/ C57BL/6 mice
K
Control (n=10)
SX 2 µg/kg (n=10)
SX 20 µg/kg (n=10)
31.5
35.5
36
Yummer 1.7 cells/ C57BL/6 mice
L
MMTV-PyMT mouse model of spontaneous breast cancer
Birth
Age (days)
Preventive treatment
Therapeutic treatment (early carcinoma)
Therapeutic treatment (late carcinoma)
Palpable tumor
Lung metastasis
M
Control (n=15)
SX 2 µg/kg (n=15)
SX 20 µg/kg (n=15)
Tumor free (%)
42
49
63
MMTV-PyMT mice
N
Control (n=15)
SX 2 µg/kg (n=15)
SX 20 µg/kg (n=15)
Number of tumor/mouse
Day 77
MMTV-PyMT mice
O
Control (n=15)
SX 20 µg/kg (n=15)
Tumor volume/mouse (mm3)
MMTV-PyMT mice
P
Control (n=15)
SX 2 µg/kg (n=15)
MMTV-PyMT mice
Q
Control (n=15)
SX 2 µg/kg (n=15)
SX 20 µg/kg (n=17)
n.s.
Day 49
MMTV-PyMT mice
R
Control (n=15)
SX 2 µg/kg (n=15)
SX 20 µg/kg (n=17)
Day 56
MMTV-PyMT mice
S
Control (n=15)
SX 20 µg/kg (n=17)
(CR, 1/17)
MMTV-PyMT mice
T
Control (n=15)
SX 2 µg/kg (n=15)
(CR, 1/15)
MMTV-PyMT mice
U
Control (n=15)
SX 2 µg/kg (n=15)
SX 20 µg/kg (n=15)
Day 70
MMTV-PyMT mice
V
Control (n=15)
SX 20 µg/kg (n=15)
MMTV-PyMT mice
W
Control (n=15)
SX 2 µg/kg (n=15)
MMTV-PyMT mice

**Figure S2**

**Figure S2. A once-weekly intraperitoneal administration of salmeterol xinafoate therapeutically inhibits the growth of established tumors and prolongs the survival of lung cancer, breast cancer, and melanoma model mice**

(**A**) Representative immunofluorescence image of a lung tissue section at day 5 from when C57BL/6 mice were isografted with LLC mouse lung cancer cells ($3 \times 10^4$) by intravenous tail vein injection. tdTomato: LLC cells, CD31: Endothelial cells, DAPI: Cell nuclei.

(**B** and **C**) Primary tumor growth curves of individual BALB/c mice orthotopically isografted with 4TO7 mouse breast cancer cells ($1 \times 10^6$). A once-weekly intraperitoneal administration of salmeterol xinafoate (20 μg/kg, B or 2 μg/kg, C) was initiated from day 8 when the primary tumor reached approximately 50 $mm^3$. The primary tumor growth curves were stopped on day 44 when the first primary tumor reached 2000 $mm^3$. Of the 20 μg/kg-treated mice, tumors disappeared in 2 of 8 mice, 1 mouse lived for at least 65 days, and 1 mouse lived for at least 59 days without detectable tumors until the experimental endpoint at day 85. The n-values denote the number of mice per group. SX: Salmeterol xinafoate. CR: Complete response.

(**D** and **G**) Primary tumor growth curves (D) and overall survival curves (G) of BALB/c mice orthotopically isografted with 4T1 mouse breast cancer cells ($1 \times 10^6$). A once-weekly intraperitoneal administration of salmeterol xinafoate (2 or 20 μg/kg) was initiated from day 7 when the primary tumor reached approximately 100 $mm^3$. The green arrow indicates the time of intraperitoneal injection of salmeterol xinafoate. The primary tumor growth curves were stopped on day 25 when the first dead mouse was observed in the control group (D). The colored number is the median survival time (G). The n-values denote the number of mice per group. SX: Salmeterol xinafoate.

(**E** and **F**) Primary tumor growth curves of individual BALB/c mice orthotopically isografted with 4T1 mouse breast cancer cells ($1 \times 10^6$). A once-weekly intraperitoneal administration of salmeterol xinafoate (20 μg/kg, E or 2 μg/kg, F) was initiated from day 7 when the primary tumor reached approximately 100 $mm^3$. The primary tumor growth curves were stopped on day 25 when the first dead mouse was observed in the control group. The n-values denote the number of mice per group. SX: Salmeterol xinafoate.

(**H** and **K**) Primary tumor growth curves (H) and overall survival curves (K) of C57BL/6 mice orthotopically isografted with Yummer 1.7 mouse melanoma cells ($3 \times 10^5$). A once-weekly intraperitoneal administration of salmeterol xinafoate (2 or 20 μg/kg) was initiated from day 10 when the primary tumor reached approximately 50 $mm^3$. The green arrow indicates the time of intraperitoneal injection of salmeterol xinafoate. The primary tumor growth curves were stopped on day 28 when the first primary tumor reached 2000 $mm^3$ (H). Mice were considered to have expired when the primary tumor reached 2000 $mm^3$ (K). The colored number is the median survival time (K). The n-values denote the number of mice per group. SX: Salmeterol xinafoate.

(**I** and **J**) Primary tumor growth curves of individual C57BL/6 mice orthotopically isografted with Yummer 1.7 mouse melanoma cells ($3 \times 10^5$). A once-weekly intraperitoneal administration of salmeterol xinafoate (20 μg/kg, I or 2 μg/kg, J) was initiated from day 10 when the primary tumor reached approximately 50 $mm^3$. The primary tumor growth curves were stopped on day 28 when the first primary tumor reached 2000 $mm^3$ in the control group. The n-values denote the number of mice per group. SX: Salmeterol xinafoate.

(**L**) Schematic of preventive treatment, treatment of early carcinoma, and treatment of late carcinoma in the MMTV-PyMT mouse model of spontaneous breast cancer. Virgin female MMTV-PyMT mice (FVB strain) spontaneously developed multiple breast tumors by approximately 42 days of age and lung metastases by approximately 67 days of age.

(**M** and **N**) Preventive treatment experiment in the MMTV-PyMT mouse model of spontaneous breast cancer (Figure S2L). The latency of spontaneous mammary tumors (M) and the number of total mammary tumors per mouse (N) in MMTV-PyMT mice with or without salmeterol xinafoate treatment on day 77. A once-weekly inhalational administration of salmeterol xinafoate (2 or 20 μg/kg) was initiated from day 28 when no palpable or visible mammary tumors existed. The colored number is the median latency time (M). The n-values denote the number of mice per group. SX: Salmeterol xinafoate.

(**O** and **P**) Preventive treatment experiment in the MMTV-PyMT mouse model of spontaneous breast cancer. Total mammary tumor growth curves of individual MMTV-PyMT mice with or without salmeterol xinafoate treatment. A once-weekly intraperitoneal administration of salmeterol xinafoate (20 μg/kg, O or 2 μg/kg, P) was initiated from day 28 when no palpable or visible mammary tumors existed. The total mammary tumor growth curves were stopped on day 77 when the first total mammary tumor per mouse reached 2000 $mm^3$ in the control group. The n-values denote the number of mice per group. SX: Salmeterol xinafoate.

(**Q**) Treatment of early carcinoma in the MMTV-PyMT mouse model of spontaneous breast cancer. Total mammary tumor volumes per mouse of MMTV-PyMT mice on day 49 before salmeterol xinafoate treatment.

(**R**) Treatment of early carcinoma in the MMTV-PyMT mouse model of spontaneous breast cancer. The number of total mammary tumors per mouse with or without salmeterol xinafoate treatment at days 56, 63, and 70. A once-weekly inhalational administration of salmeterol xinafoate (2 or 20 μg/kg) was initiated on day 49 when palpable or visible mammary tumors existed. The n-values denote the number of mice per group. SX: Salmeterol xinafoate.

(**S** and **T**) Treatment of early carcinoma in the MMTV-PyMT mouse model of spontaneous breast cancer. Total mammary tumor growth curves of individual MMTV-PyMT mice with or without salmeterol xinafoate treatment. A once-weekly intraperitoneal administration of salmeterol xinafoate (20 μg/kg, S or 2 μg/kg, T) was initiated from day 49 when palpable or visible mammary tumors existed. The total mammary tumor growth curves were stopped on day 84 when the first total mammary tumor per mouse reached 2000 $mm^3$ in the control group. Of the 20 μg/kg-treated mice, tumors disappeared in 1 of 17 mice, and the mice lived for at least 47 days without detectable tumors until the experimental endpoint at day 180. Of the 2 μg/kg-treated mice, tumors disappeared in 1 of 15 mice, and the mice lived for at least 96 days without detectable tumors until the experimental endpoint at day 180. The n-values denote the number of mice per group. SX: Salmeterol xinafoate. CR: Complete response.

(**U**) Treatment of late carcinoma in the MMTV-PyMT mouse model of spontaneous breast cancer. Total mammary tumor volumes per mouse of MMTV-PyMT mice on day 70 before salmeterol xinafoate treatment.

(**V** and **W**) Treatment of late carcinoma in the MMTV-PyMT mouse model of spontaneous breast cancer. Total mammary tumor growth curves of individual MMTV-PyMT mice with or without salmeterol xinafoate treatment. A once-weekly intraperitoneal administration of salmeterol xinafoate (20 μg/kg, V or 2 μg/kg, W) was initiated from day 70 when the total mammary tumor reached approximately 800 $mm^3$ and lung metastasis developed spontaneously. The total mammary tumor growth curves were stopped on day 80 when the first total mammary tumor per mouse reached 2000 $mm^3$ in the control group. The n-values denote the number of mice per group. SX: Salmeterol xinafoate.

Anesthetics were not used in the mouse experiments, except for cancer cells injection, because anesthetics prevented the anticancer effects of salmeterol xinafoate. Mice were fasted for 12 h before salmeterol xinafoate administration. Eight hours after salmeterol xinafoate administration, food was provided. The data are presented as the mean ± s.e.m. values. *P*-values were determined by unpaired two-way ANOVA with uncorrected Fisher's LSD test (D, H, and R), unpaired one-way ANOVA with uncorrected Fisher's LSD test (N, Q, and U), or the log-rank test (G, K, and M). $*P < 0.05$, $**P < 0.01$, $***P < 0.001$, $****P < 0.0001$, and n.s., not significant.

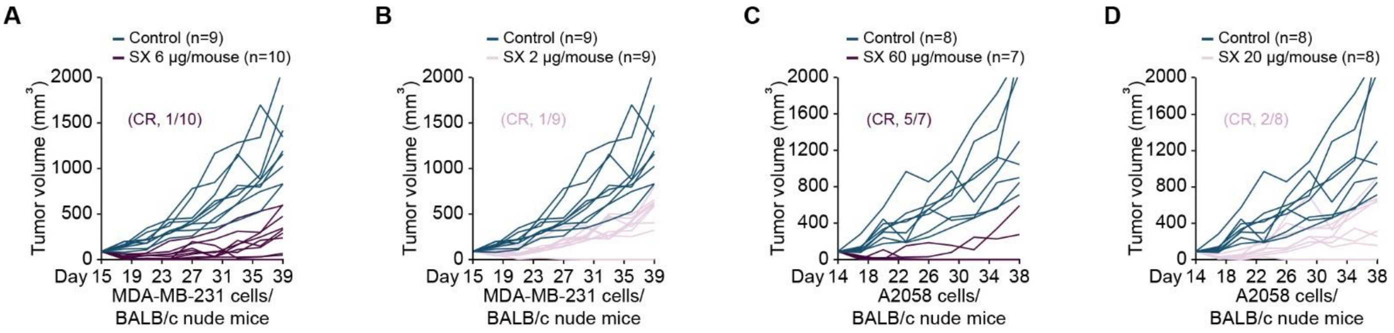


Figure S3

**Figure S3. A once-weekly inhalational or transdermal administration of salmeterol xinafoate therapeutically inhibits the growth of established tumors and prolongs survival of model mice with human breast cancer and melanoma**

(**A** and **B**) Primary tumor growth curves of individual BALB/c nude mice orthotopically xenografted with MDA-MB-231 human breast cancer cells ($1 \times 10^6$). A once-weekly inhalational administration of salmeterol xinafoate (6 μg/mouse, A or 2 μg/mouse, B) was initiated from day 15 when the primary tumor reached approximately 90 $mm^3$. The primary tumor growth curves were stopped on day 39 when the first primary tumor reached 2000 $mm^3$ in the control group. Of the 6 μg/mouse-treated mice, tumors disappeared in 1 of 10 mice, and the mice lived for at least 63 days without detectable tumor until the experimental endpoint at day 90. Of the 2 μg/mouse-treated mice, tumors disappeared in 1 of 9 mice, and the mice lived for at least 60 days without detectable tumor until the experimental endpoint at day 90. The n-values denote the number of mice per group. SX: Salmeterol xinafoate. CR: Complete response.

(**C** and **D**) Primary tumor growth curves of individual BALB/c nude mice orthotopically xenografted with A2058 human melanoma cells ($1 \times 10^6$). A once-weekly transdermal administration of salmeterol xinafoate (60 μg/mouse, C or 20 μg/mouse, D) was initiated from day 14 when the primary tumor reached approximately 100 $mm^3$. The primary tumor growth curves were stopped on day 38 when the first primary tumor reached 2000 $mm^3$ in the control group. Of the 60 μg/mouse-treated mice, melanoma disappeared in 5 of 7 mice, 4 mice lived for at least 152 days, and 1 mouse lived for at least 149 days without detectable tumors until the time of manuscript submission on day 175. Of the 20 μg/mouse-treated mice, melanoma disappeared in 2 of 8 mice, 1 mouse lived for at least 149 days, and 1 mouse lived for at least 143 days without detectable tumors until the time of manuscript submission on day 175. The n-values denote the number of mice per group. SX: Salmeterol xinafoate. CR: Complete response.

Anesthetics were not used in the mouse experiments, except for cancer cells injection, because anesthetics prevented the anticancer effects of salmeterol xinafoate. Mice were fasted for 6 h before salmeterol xinafoate administration. Four hours after salmeterol xinafoate administration, food was provided.

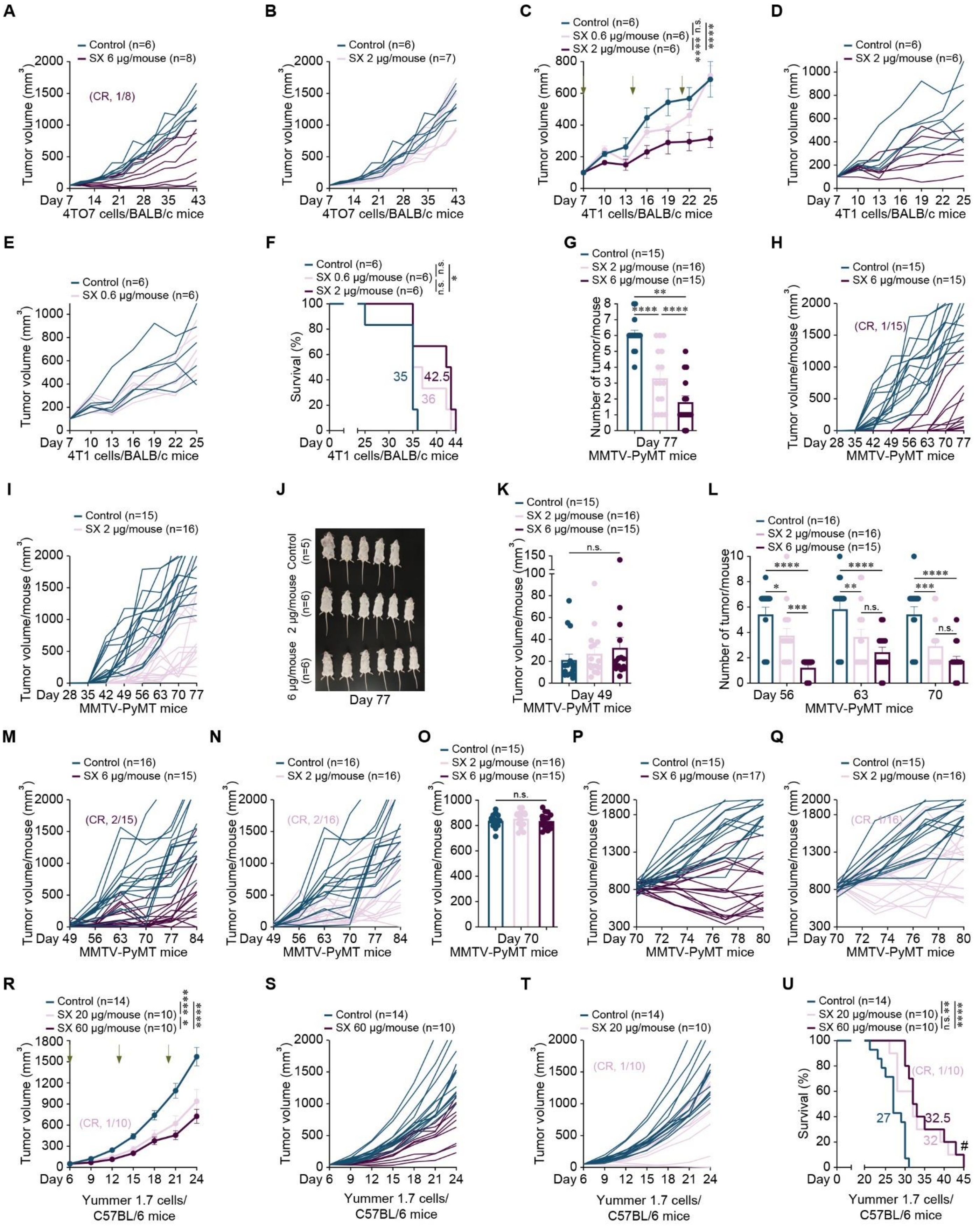

A
Control (n=6)
SX 6 μg/mouse (n=8)
(CR, 1/8)
Tumor volume (mm3)
4TO7 cells/BALB/c mice
B
Control (n=6)
SX 2 μg/mouse (n=7)
C
Control (n=6)
SX 0.6 μg/mouse (n=6)
SX 2 μg/mouse (n=6)
4T1 cells/BALB/c mice
D
Control (n=6)
SX 2 μg/mouse (n=6)
E
Control (n=6)
SX 0.6 μg/mouse (n=6)
F
Control (n=6)
SX 0.6 μg/mouse (n=6)
SX 2 μg/mouse (n=6)
Survival (%)
G
Control (n=15)
SX 2 μg/mouse (n=16)
SX 6 μg/mouse (n=15)
Number of tumor/mouse
Day 77
MMTV-PyMT mice
H
Control (n=15)
SX 6 μg/mouse (n=15)
(CR, 1/15)
Tumor volume/mouse (mm3)
I
Control (n=15)
SX 2 μg/mouse (n=16)
J
Control (n=5)
2 μg/mouse (n=6)
6 μg/mouse (n=6)
Day 77
K
Control (n=15)
SX 2 μg/mouse (n=16)
SX 6 μg/mouse (n=15)
Day 49
L
Control (n=16)
SX 2 μg/mouse (n=16)
SX 6 μg/mouse (n=15)
M
Control (n=16)
SX 6 μg/mouse (n=15)
(CR, 2/15)
N
Control (n=16)
SX 2 μg/mouse (n=16)
(CR, 2/16)
O
Control (n=15)
SX 2 μg/mouse (n=16)
SX 6 μg/mouse (n=15)
Day 70
P
Control (n=15)
SX 6 μg/mouse (n=17)
Q
Control (n=15)
SX 2 μg/mouse (n=16)
(CR, 1/16)
R
Control (n=14)
SX 20 μg/mouse (n=10)
SX 60 μg/mouse (n=10)
(CR, 1/10)
Yummer 1.7 cells/
C57BL/6 mice
S
Control (n=14)
SX 60 μg/mouse (n=10)
T
Control (n=14)
SX 20 μg/mouse (n=10)
(CR, 1/10)
U
Control (n=14)
SX 20 μg/mouse (n=10)
SX 60 μg/mouse (n=10)
(CR, 1/10)


**Figure S4**

**Figure S4. A once-weekly inhalational or transdermal administration of salmeterol xinafoate therapeutically inhibits the growth of established tumors and prolongs the survival of breast cancer and melanoma model mice**

(**A** and **B**) Primary tumor growth curves of individual BALB/c mice orthotopically isografted with 4TO7 mouse breast cancer cells ($1 \times 10^6$). A once-weekly inhalational administration of salmeterol xinafoate (6 μg/mouse, A or 2 μg/mouse, B) was initiated from day 7 when the primary tumor reached approximately 50 $mm^3$. The primary tumor growth curves were stopped on day 43 when the first primary tumor reached 2000 $mm^3$ in the control group. Of the 6 μg/mouse-treated mice, tumors disappeared in 1 of 8 mice, and the mice lived for at least 44 days without detectable tumors until the experimental endpoint at day 96 (A). The n-values denote the number of mice per group. SX: Salmeterol xinafoate. CR: Complete response.

(**C** and **F**) Primary tumor growth curves (C) and overall survival curves (F) of BALB/c mice orthotopically isografted with 4T1 mouse breast cancer cells ($1 \times 10^6$). A once-weekly inhalational administration of salmeterol xinafoate (0.6 or 2 μg/mouse) was initiated from day 7 when the primary tumor reached approximately 100 $mm^3$. The green arrow indicates the time of inhalational administration of salmeterol xinafoate. The primary tumor growth curves were stopped on day 25 when the first dead mouse was observed in the control group (C). The colored number is the median survival time (F). The n-values denote the number of mice per group. SX: Salmeterol xinafoate.

(**D** and **E**) Primary tumor growth curves of individual BALB/c mice orthotopically isografted with 4T1 mouse breast cancer cells ($1 \times 10^6$). A once-weekly inhalational administration of salmeterol xinafoate (2 μg/mouse, D or 0.6 μg/mouse, E) was initiated from day 7 when the primary tumor reached approximately 100 $mm^3$. The primary tumor growth curves were stopped on day 25 when the first dead mouse was observed in the control group. The n-values denote the number of mice per group. SX: Salmeterol xinafoate.

(**G**) Preventive treatment experiment in the MMTV-PyMT mouse model of spontaneous breast cancer. The number of total mammary tumors per mouse with or without salmeterol xinafoate treatment at day 77. A once-weekly inhalational administration of salmeterol xinafoate (2 or 6 μg/mouse) was initiated from day 28 when no palpable or visible mammary tumors existed. The n-values denote the number of mice per group. SX: Salmeterol xinafoate.

(**H** and **I**) Preventive treatment experiment in the MMTV-PyMT mouse model of spontaneous breast cancer. Total mammary tumor growth curves of individual MMTV-PyMT mice with or without salmeterol xinafoate treatment. A once-weekly inhalational administration of salmeterol xinafoate (6 μg/mouse, H or 2 μg/mouse, I) was initiated from day 28 when no palpable or visible mammary tumors existed. The total mammary tumor growth curves were stopped on day 77 when the first total mammary tumor per mouse reached 2000 $mm^3$ in the control group. Of the 6 μg/mouse-treated mice, tumors disappeared in 1 of 15 mice, and the mice lived for at least 61 days without detectable tumors until the experimental endpoint at day 180 (H). The n-values denote the number of mice per group. SX: Salmeterol xinafoate. CR: Complete response.

(**J**) Independent preventive treatment experiment in the MMTV-PyMT mouse model of spontaneous breast cancer showing the image of individual MMTV-PyMT mice with or without salmeterol xinafoate treatment at day 77. A once-weekly inhalational administration of salmeterol xinafoate (2 or 6 μg/mouse) was initiated from day 28 when no palpable or visible mammary tumors existed. The n-values denote the number of mice per group. SX: Salmeterol xinafoate.

(**K**) Treatment of early carcinoma in the MMTV-PyMT mouse model of spontaneous breast cancer. Total mammary tumor volumes per mouse of MMTV-PyMT mice on day 49 before salmeterol xinafoate treatment.

(**L**) Treatment of early carcinoma in the MMTV-PyMT mouse model of spontaneous breast cancer. The number of total mammary tumors per mouse with or without salmeterol xinafoate treatment at days 56, 63, and 70. A once-weekly inhalational administration of salmeterol xinafoate (2 or 6 μg/mouse) was initiated on day 49 when palpable or visible mammary tumors existed. The n-values denote the number of mice per group. SX: Salmeterol xinafoate.

(**M** and **N**) Treatment of early carcinoma in the MMTV-PyMT mouse model of spontaneous breast cancer. Total mammary tumor growth curves of individual MMTV-PyMT mice with or without salmeterol xinafoate treatment. A once-weekly inhalational administration of salmeterol xinafoate (6 μg/mouse, M or 2 μg/mouse, N) was initiated from day 49 when palpable or visible mammary tumors existed. The total mammary tumor growth curves were stopped on day 84 when the first total mammary tumor per mouse reached 2000 $mm^3$ in the control group. Of the 6 μg/mouse-treated mice, tumors disappeared in 2 of 15 mice, 1 mouse lived for at least 96 days, and 1 mouse lived for at least 82 days without detectable tumors until the experimental endpoint at day 180 (M). Of the 2 μg/mouse-treated mice, tumors disappeared in 2 of 16 mice, and the mice lived for at least 68 days without detectable tumors until the experimental endpoint at day 180 (N). The n-values denote the number of mice per group. SX: Salmeterol xinafoate. CR: Complete response.

(**O**) Treatment of late carcinoma in the MMTV-PyMT mouse model of spontaneous breast cancer. Total mammary tumor volumes per mouse of MMTV-PyMT mice on day 70 before salmeterol xinafoate treatment.

(**P** and **Q**) Treatment of late carcinoma in the MMTV-PyMT mouse model of spontaneous breast cancer. Total mammary tumor growth curves of individual MMTV-PyMT mice with or without salmeterol xinafoate treatment. A once-weekly inhalational administration of salmeterol xinafoate (6 μg/mouse, P or 2 μg/mouse, Q) was initiated from day 70 when the total mammary tumor reached approximately 800 $mm^3$ and lung metastasis developed spontaneously. The total mammary tumor growth curves were stopped on day 80 when the first total mammary tumor per mouse reached 2000 $mm^3$ in the control group. Of the 2 μg/mouse-treated mice, tumors disappeared in 1 of 16 mice, and the mice lived for at least 54 days without detectable tumors until the experimental endpoint at day 180 (Q). The n-values denote the number of mice per group. SX: Salmeterol xinafoate. CR: Complete response.

(**R** and **U**) Primary tumor growth curves (R) and overall survival curves (U) of C57BL/6 mice orthotopically isografted with Yummer 1.7 mouse melanoma cells ($3 \times 10^5$). A once-weekly transdermal administration of salmeterol xinafoate (20 or 60 μg/mouse) was initiated from day 6 when the primary tumor reached approximately 50 $mm^3$. The green arrow indicates the time of inhalational administration of salmeterol xinafoate. The primary tumor growth curves were stopped on day 24 when the first primary tumor reached 2000 $mm^3$ in the control group (R). Mice were considered to have expired when the primary tumor reached 2000 $mm^3$ (U). Of the 20 μg/mouse-treated mice, tumors disappeared in 1 of 10 mice, and the mice lived for at least 116 days without detectable tumors until the experimental endpoint at day 140. The colored number is the median survival time (U). The n-values denote the number of mice per group. SX: Salmeterol xinafoate. # The experimental endpoint on day 140.

(**S** and **T**) Primary tumor growth curves of individual C57BL/6 mice orthotopically isografted with Yummer 1.7 mouse melanoma cells ($3 \times 10^5$). A once-weekly transdermal administration of salmeterol xinafoate (60 μg/mouse, S or 20 μg/mouse, T) was initiated from day 6 when the primary tumor reached approximately 50 $mm^3$. The primary tumor growth curves were stopped on day 24 when the first primary tumor reached 2000 $mm^3$ in the control group. Of the 20 μg/mouse-treated mice, tumors disappeared in 1 of 10 mice, and the mice lived for at least 116 days without detectable tumors until the experimental endpoint at day 140 (T). The n-values denote the number of mice per group. SX: Salmeterol xinafoate.

Anesthetics were not used in the mouse experiments, except for cancer cells injection, because anesthetics prevented the anticancer effects of salmeterol xinafoate. Mice were fasted for 12 h before salmeterol xinafoate administration. Eight hours after salmeterol xinafoate administration, food was provided. The data are presented as the mean ± s.e.m. values. *P*-values were determined by unpaired two-way ANOVA with uncorrected Fisher's LSD test (C, L, and R), unpaired one-way ANOVA with uncorrected Fisher's

LSD test (G, K, and O), or the log-rank test (F and U). *$P < 0.05$, **$P < 0.01$, ***$P < 0.001$, ****$P < 0.0001$, and n.s., not significant.

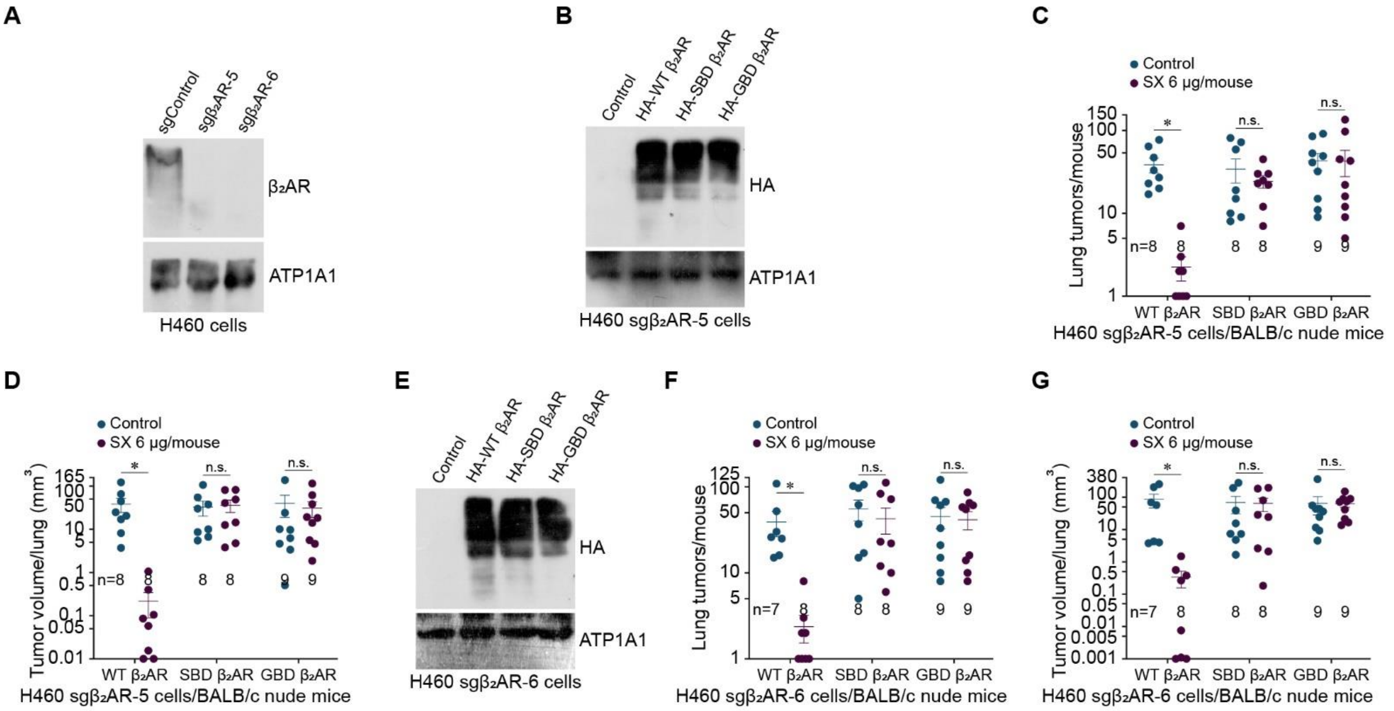


Figure S5

**Figure S5. Salmeterol xinafoate exerts anticancer effects through β2AR expressed on cancer cells and is dependent on the downstream signaling pathway in a mouse model of human lung cancer**

(**A**) Immunoblot analysis of endogenous β2AR expression on the cell membrane of control or β2AR-knockout H460 lung cancer cells. ATP1A1: Cell membrane protein (three independent experiments).

(**B** and **E**) Immunoblot analysis of membrane expression of control, exogenous HA-tagged wild-type human β2AR (HA-WT β2AR), salmeterol xinafoate-binding-deficient human β2AR (F193W, F194V, S203T, S204T, H296K, R304P, K305D, and Y308F mutations, HA-SBD β2AR), or G-protein-binding-deficient human β2AR (R131A and F139A mutations, HA-GBD β2AR) in endogenous β2AR-knockout H460 lung cancer cells (sg$\beta_2$AR-5, B and sg$\beta_2$AR-6, E). ATP1A1: Cell membrane protein (three independent experiments).

(**C** and **D**) The number of tumors per lung (C) and the tumor volume per lung (D) on day 51 from when BALB/c nude mice were xenografted with H460 human lung cancer cells ($1 \times 10^6$) by intravenous tail vein injection. In endogenous $\beta_2$AR knockout H460 lung cancer cells (sg$\beta_2$AR-5), wild-type human $\beta_2$AR (WT $\beta_2$AR), salmeterol xinafoate-binding-deficient human $\beta_2$AR (F193W, F194V, S203T, S204T, H296K, R304P, K305D, and Y308F mutations, SBD $\beta_2$AR), or G-protein-binding-deficient human $\beta_2$AR (R131A and F139A mutations, GBD $\beta_2$AR) was overexpressed as indicated. A once-weekly inhalational administration of salmeterol xinafoate (6 μg/mouse) was initiated on day 21 from when the lung tumor was detected. The experimental endpoint was on day 51 when the first dead mouse was observed in the control group. All lungs were analyzed by serial sectioning the entire lung and scoring tumor nodules. A tumor nodule larger than 100 μm in width was counted and calculated as described in the Materials and Methods. Of the 6 μg/mouse-treated mice, which were xenografted with the indicated H460 sg$\beta_2$AR-5/WT $\beta_2$AR cells, lung tumors disappeared in 2 of 8 mice. The n-values denote the number of mice per group. SX: Salmeterol xinafoate.

(**F** and **G**) The number of tumors per lung (F) and the tumor volume per lung (G) on day 51 from BALB/c nude mice xenografted with H460 human lung cancer cells (1 × $10^6$) by intravenous tail vein injection. In endogenous $\beta_2$AR knockout H460 lung cancer cells (sg$\beta_2$AR-6), wild-type human $\beta_2$AR (WT $\beta_2$AR), salmeterol xinafoate-binding-deficient human $\beta_2$AR (F193W, F194V, S203T, S204T, H296K, R304P, K305D, and Y308F mutations, SBD $\beta_2$AR), or G-protein-binding-deficient human $\beta_2$AR (R131A and F139A mutations, GBD $\beta_2$AR) was overexpressed as indicated. A once-weekly inhalational administration of salmeterol xinafoate (6 μg/mouse) was initiated on day 21 from when the lung tumor was detected. The experimental endpoint was on day 51 when the first dead mouse was observed in the control group. All lungs were analyzed by serial sectioning the entire lung and scoring tumor nodules. A tumor nodule larger than 100 μm in width was counted and calculated as described in the Materials and Methods. Of the 6 μg/mouse-treated mice, which were xenografted with the indicated H460 sg$\beta_2$AR-6/WT $\beta_2$AR cells, lung tumors disappeared in 2 of 8 mice. The n-values denote the number of mice per group. SX: Salmeterol xinafoate.

Anesthetics were not used in the mouse experiments, except for cancer cells injection, because anesthetics prevented the anticancer effects of salmeterol xinafoate. Mice were fasted for 6 h before salmeterol xinafoate administration. Four hours after salmeterol xinafoate administration, food was provided. The data are presented as the mean ± s.e.m.

values. *P*-values were determined by unpaired two-way ANOVA with uncorrected Fisher's LSD test (C, D, F, and G). *$P < 0.05$ and n.s., not significant.

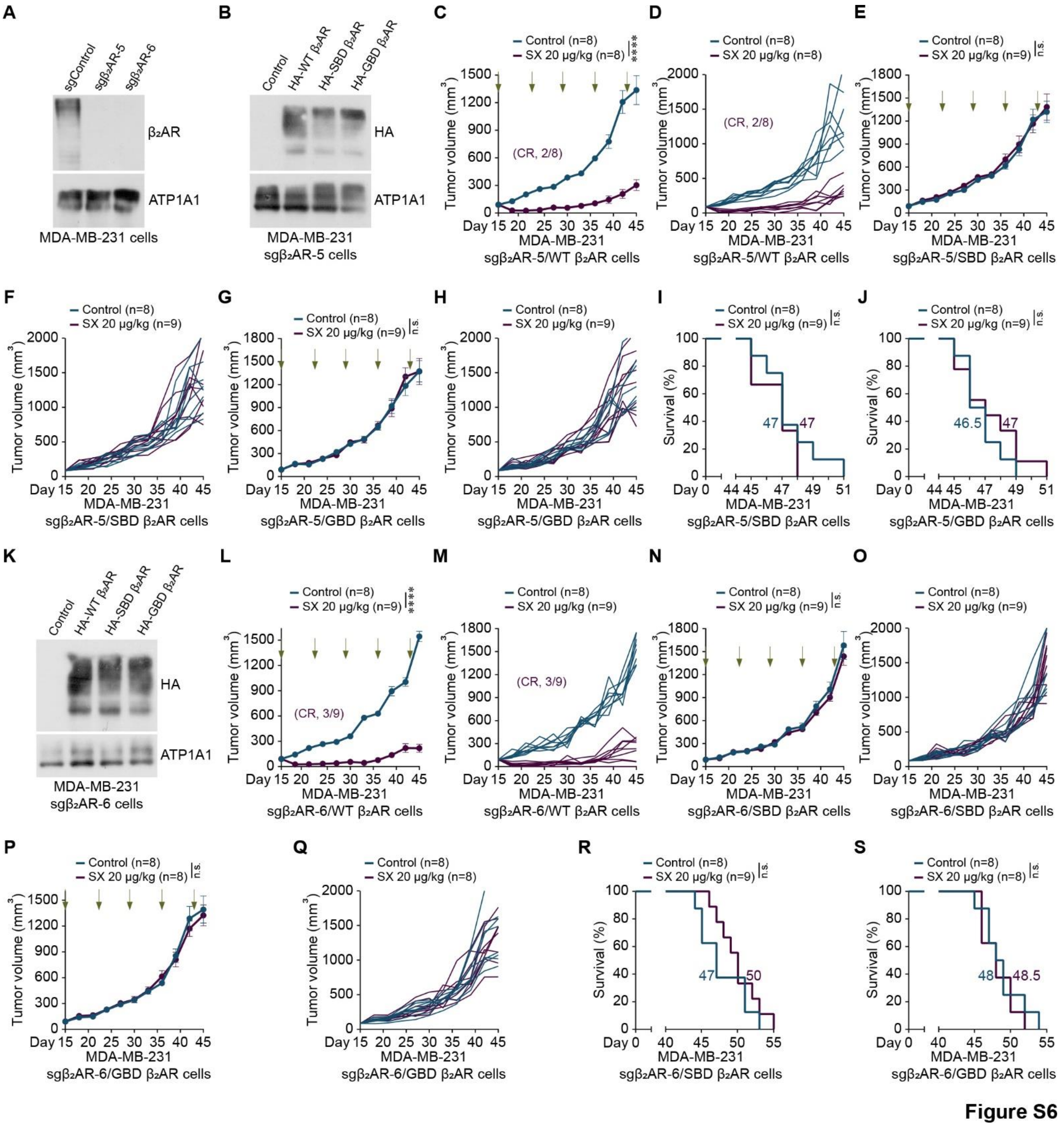


**Figure S6**

**Figure S6. Salmeterol xinafoate exerts anticancer effects through β2AR expressed on cancer cells and is dependent on the downstream signaling pathway in a mouse model of human breast cancer**

(**A**) Immunoblot analysis of endogenous $\beta_2$AR expression on the cell membrane of control or $\beta_2$AR-knockout MDA-MB-231 breast cancer cells. ATP1A1: Cell membrane protein (three independent experiments).

(**B** and **K**) Immunoblot analysis of membrane expression of control, exogenous HA-tagged wild-type human $\beta_2$AR (HA-WT $\beta_2$AR), salmeterol xinafoate-binding-deficient human $\beta_2$AR (F193W, F194V, S203T, S204T, H296K, R304P, K305D, and Y308F mutations, HA-SBD $\beta_2$AR), or G-protein-binding-deficient human $\beta_2$AR (R131A and F139A mutations, HA-GBD $\beta_2$AR) in endogenous $\beta_2$AR-knockout MDA-MB-231 breast cancer cells (sg$\beta_2$AR-5, B and sg$\beta_2$AR-6, K). ATP1A1: Cell membrane protein (three independent experiments).

(**C** and **D**) Average primary tumor growth curves (C) and individual primary tumor growth curves (D) of BALB/c nude mice orthotopically xenografted with the indicated MDA-MB-231 human breast cancer cells ($1 \times 10^6$), which overexpressed the wild-type human $\beta_2$AR (WT $\beta_2$AR) in endogenous $\beta_2$AR knockout MDA-MB-231 breast cancer cells (sg$\beta_2$AR-5). A once-weekly intraperitoneal administration of salmeterol xinafoate (20 μg/kg) was initiated from day 15 when the primary tumor reached approximately 90 $mm^3$. The green arrow indicates the time of intraperitoneal injection of salmeterol xinafoate. The primary tumor growth curves were stopped on day 45 when the first primary tumor reached 2000 $mm^3$ in the control group. Of the 20 μg/kg-treated mice, tumors disappeared in 2 of 8 mice, and the 2 mice lived for at least 139 and 124 days without detectable tumors until the experimental endpoint at day 172. The n-values denote the number of mice per group. SX: Salmeterol xinafoate. CR: Complete response.

(**E**, **F**, and **I**) Average primary tumor growth curves (E), individual primary tumor growth curves (F), and overall survival curves (I) of BALB/c nude mice orthotopically xenografted with the indicated MDA-MB-231 human breast cancer cells ($1 \times 10^6$), which overexpressed the salmeterol xinafoate-binding-deficient human $\beta_2$AR (F193W, F194V, S203T, S204T, H296K, R304P, K305D, and Y308F mutations, SBD $\beta_2$AR) in endogenous $\beta_2$AR knockout MDA-MB-231 breast cancer cells (sg$\beta_2$AR-5). A once-weekly intraperitoneal administration of salmeterol xinafoate (20 μg/kg) was initiated from day 15 when the primary tumor reached approximately 90 $mm^3$. The green arrow indicates the time of intraperitoneal injection of salmeterol xinafoate. The primary tumor growth curves were stopped on day 45 when the first primary tumor reached 2000 $mm^3$ in the control group (E and F). Mice were considered to have expired when the primary tumor reached 2000 $mm^3$ (I). The colored number is the median survival time (I). The n-values denote the number of mice per group. SX: Salmeterol xinafoate.

(**G**, **H**, and **J**) Average primary tumor growth curves (G), individual primary tumor growth curves (H), and overall survival curves (J) of BALB/c nude mice orthotopically xenografted with the indicated MDA-MB-231 human breast cancer cells ($1 \times 10^6$), which overexpressed the G-protein-binding-deficient human $\beta_2$AR (R131A and F139A mutations, GBD $\beta_2$AR) in endogenous $\beta_2$AR knockout MDA-MB-231 breast cancer cells (sg$\beta_2$AR-5). A once-weekly intraperitoneal administration of salmeterol xinafoate (20 μg/kg) was initiated from day 15 when the primary tumor reached approximately 90 $mm^3$. The green arrow indicates the time of intraperitoneal injection of salmeterol xinafoate. The primary tumor growth curves were stopped on day 45 when the first primary tumor reached 2000 $mm^3$ in the control group (G and H). Mice were considered to have expired when the primary tumor reached 2000 $mm^3$ (J). The colored number is the median survival time (J). The n-values denote the number of mice per group. SX: Salmeterol xinafoate.

(**L** and **M**) Average primary tumor growth curves (L) and individual primary tumor growth curves (M) of BALB/c nude mice orthotopically xenografted with the indicated MDA-MB-231 human breast cancer cells ($1 \times 10^6$), which overexpressed the wild-type human $\beta_2$AR (WT $\beta_2$AR) in endogenous $\beta_2$AR knockout MDA-MB-231 breast cancer cells (sg$\beta_2$AR-6). A once-weekly intraperitoneal administration of salmeterol xinafoate (20 μg/kg) was initiated from day 15 when the primary tumor reached approximately 90 $mm^3$. The green arrow indicates the time of intraperitoneal injection of salmeterol xinafoate. The primary tumor growth curves were stopped on day 45 when the first primary tumor reached 2000 $mm^3$ in the control group. Of the 20 μg/kg-treated mice, tumors disappeared in 3 of 9 mice, and the 3 mice lived for at least 175, 148, and 139 days, without detectable tumors until the experimental endpoint at day 199. The n-values denote the number of mice per group. SX: Salmeterol xinafoate. CR: Complete response.

(**N**, **O**, and **R**) Average primary tumor growth curves (N), individual primary tumor growth curves (O), and overall survival curves (R) of BALB/c nude mice orthotopically xenografted with the indicated MDA-MB-231 human breast cancer cells ($1 \times 10^6$), which overexpressed the salmeterol xinafoate-binding-deficient human $\beta_2$AR (F193W, F194V, S203T, S204T, H296K, R304P, K305D, and Y308F mutations, SBD $\beta_2$AR) in endogenous $\beta_2$AR knockout MDA-MB-231 breast cancer cells (sg$\beta_2$AR-6). A once-weekly intraperitoneal administration of salmeterol xinafoate (20 μg/kg) was initiated from day 15 when the primary tumor reached approximately 90 $mm^3$. The green arrow indicates the time of intraperitoneal injection of salmeterol xinafoate. The primary tumor growth curves were stopped on day 45 when the first primary tumor reached 2000 $mm^3$ in the control group (N and O). Mice were considered to have expired when the primary tumor reached 2000 $mm^3$ (R). The colored number is the median survival time (R). The n-values denote the number of mice per group. SX: Salmeterol xinafoate.

(**P**, **Q**, and **S**) Average primary tumor growth curves (P), individual primary tumor growth curves (Q), and overall survival curves (S) of BALB/c nude mice orthotopically xenografted with the indicated MDA-MB-231 human breast cancer cells ($1 \times 10^6$), which overexpressed the G-protein-binding-deficient human $\beta_2$AR (R131A and F139A mutations, GBD $\beta_2$AR) in endogenous $\beta_2$AR knockout MDA-MB-231 breast cancer cells (sg$\beta_2$AR-6). A once-weekly intraperitoneal administration of salmeterol xinafoate (20 μg/kg) was initiated from day 15 when the primary tumor reached approximately 90 $mm^3$. The green arrow indicates the time of intraperitoneal injection of salmeterol xinafoate. The primary tumor growth curves were stopped on day 45 when the first primary tumor reached 2000 $mm^3$ in the control group (P and Q). Mice were considered to have expired when the primary tumor reached 2000 $mm^3$ (S). The colored number is the median survival time (S). The n-values denote the number of mice per group. SX: Salmeterol xinafoate.

Anesthetics were not used in the mouse experiments, except for cancer cells injection, because anesthetics prevented the anticancer effects of salmeterol xinafoate. Mice were fasted for 6 h before salmeterol xinafoate administration. Four hours after salmeterol xinafoate administration, food was provided. The data are presented as the mean ± s.e.m. values. *P*-values were determined by unpaired two-way ANOVA with uncorrected Fisher's LSD test (C, E, G, L, N, and P) or the log-rank test (I, J, R, and S). $^{****}P < 0.0001$ and n.s., not significant.

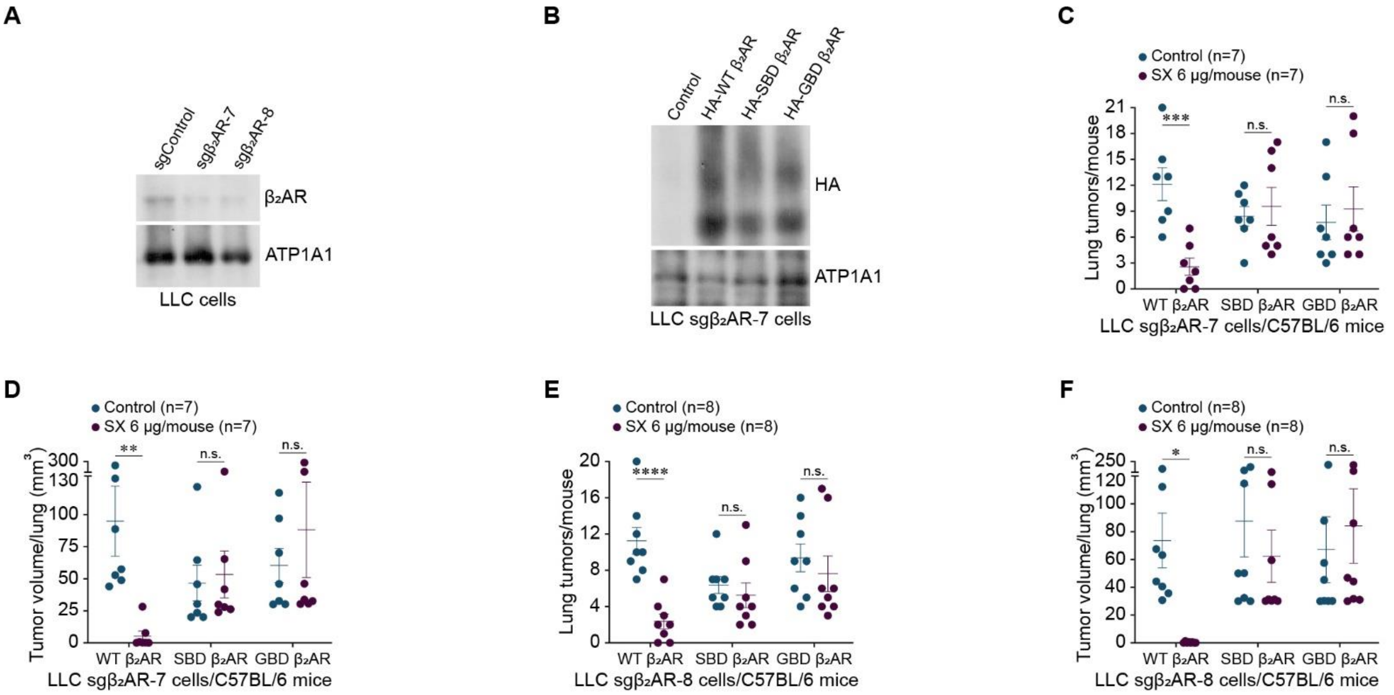


**Figure S7**

**Figure S7. Salmeterol xinafoate exerts anticancer effects through β2AR expressed on cancer cells and is dependent on the downstream signaling pathway in lung cancer model mice**

(**A**) Immunoblot analysis of endogenous β2AR expression on the cell membrane of control or β2AR-knockout LLC lung cancer cells. ATP1A1: Cell membrane protein (three independent experiments).

(**B**) Immunoblot analysis of membrane expression of control, exogenous HA-tagged wild-type mouse β2AR (HA-WT β2AR), salmeterol xinafoate-binding-deficient mouse β2AR (F193W, F194V, S203T, S204T, H296K, K305D, and Y308F mutations, HA-SBD β2AR), or G-protein-binding-deficient mouse β2AR (R131A and F139A mutations, HA-GBD $\beta_2$AR) in endogenous $\beta_2$AR-knockout LLC lung cancer cells (sg$\beta_2$AR-7). ATP1A1: Cell membrane protein (three independent experiments).

(**C** and **D**) The number of tumors per lung (C) and the tumor volume per lung (D) on day 39 from when C57BL/6 mice were isografted with LLC mouse lung cancer cells ($3 \times 10^4$) by intravenous tail vein injection. In endogenous $\beta_2$AR knockout LLC lung cancer cells (sg$\beta_2$AR-7), wild-type mouse $\beta_2$AR (WT $\beta_2$AR), salmeterol xinafoate-binding-deficient mouse $\beta_2$AR (F193W, F194V, S203T, S204T, H296K, K305D, and Y308F mutations, SBD $\beta_2$AR), or G-protein-binding-deficient mouse $\beta_2$AR (R131A and F139A mutations, GBD $\beta_2$AR) was overexpressed as indicated. A once-weekly inhalational administration of salmeterol xinafoate (6 μg/mouse) was initiated on day 5 from when the lung tumor was detected. The experimental endpoint was on day 39 when the first dead mouse was observed in the control group. All lungs were analyzed by serial sectioning the entire lung and scoring tumor nodules. A tumor nodule larger than 100 μm in width was counted and calculated as described in the Materials and Methods. Of the 6 μg/mouse-treated mice, which were isografted with the indicated LLC sg$\beta_2$AR-7/WT $\beta_2$AR cells, lung tumors disappeared in 2 of 8 mice. The n-values denote the number of mice per group. SX: Salmeterol xinafoate.

(**E** and **F**) The number of tumors per lung (E) and the tumor volume per lung (F) on day 39 from when C57BL/6 mice were isografted with LLC mouse lung cancer cells

($3 \times 10^4$) by intravenous tail vein injection. In endogenous $\beta_2$AR knockout LLC lung cancer cells (sg$\beta_2$AR-8), wild-type mouse $\beta_2$AR (WT $\beta_2$AR), salmeterol xinafoate-binding-deficient mouse $\beta_2$AR (F193W, F194V, S203T, S204T, H296K, K305D, and Y308F mutations, SBD $\beta_2$AR), or G-protein-binding-deficient mouse $\beta_2$AR (R131A and F139A mutations, GBD $\beta_2$AR) was overexpressed as indicated. A once-weekly inhalational administration of salmeterol xinafoate (6 μg/mouse) was initiated on day 5 from when the lung tumor was detected. The experimental endpoint was on day 39 when the first dead mouse was observed in the control group. All lungs were analyzed by serial sectioning the entire lung and scoring tumor nodules. A tumor nodule larger than 100 μm in width was counted and calculated as described in the Materials and Methods. Of the 6 μg/mouse-treated mice, which were isografted with the indicated LLC sg$\beta_2$AR-8/WT $\beta_2$AR cells, lung tumors disappeared in 2 of 8 mice. The n-values denote the number of mice per group. SX: Salmeterol xinafoate.

Anesthetics were not used in the mouse experiments, except for cancer cells injection, because anesthetics prevented the anticancer effects of salmeterol xinafoate. Mice were fasted for 12 h before salmeterol xinafoate administration. Eight hours after salmeterol xinafoate administration, food was provided. The data are presented as the mean ± s.e.m. values. *P*-values were determined by unpaired two-way ANOVA with uncorrected Fisher's LSD test (C-F). *$P < 0.05$, **$P < 0.01$, ***$P < 0.001$, and ****$P < 0.0001$, and n.s., not significant.

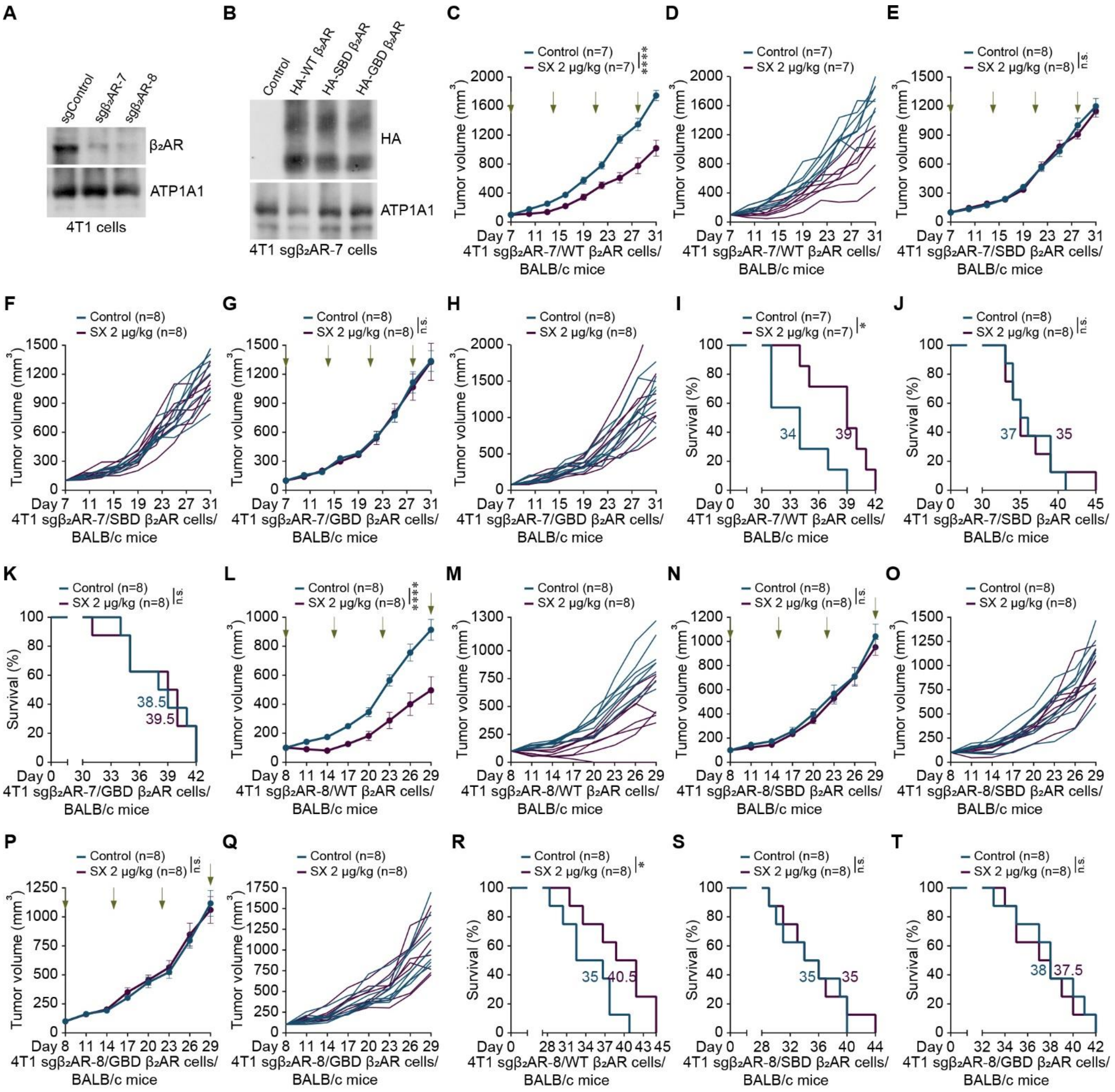


**Figure S8**

**Figure S8. Salmeterol xinafoate exerts anticancer effects through β2AR expressed on cancer cells and is dependent on the downstream signaling pathway in breast cancer model mice**

(**A**) Immunoblot analysis of endogenous $\beta_2$AR expression on the cell membrane of control or $\beta_2$AR-knockout 4T1 breast cancer cells. ATP1A1: Cell membrane protein (three independent experiments).

(**B**) Immunoblot analysis of membrane expression of control, exogenous HA-tagged wild-type mouse $\beta_2$AR (HA-WT $\beta_2$AR), salmeterol xinafoate-binding-deficient mouse $\beta_2$AR (F193W, F194V, S203T, S204T, H296K, K305D, and Y308F mutations, HA-SBD $\beta_2$AR), or G-protein-binding-deficient mouse $\beta_2$AR (R131A and F139A mutations, HA-GBD $\beta_2$AR) in endogenous $\beta_2$AR-knockout 4T1 breast cancer cells (sg$\beta_2$AR-7). ATP1A1: Cell membrane protein (three independent experiments).

(**C**, **D**, and **I**) Average primary tumor growth curves (C), individual primary tumor growth curves (D), and overall survival curves (I) of BALB/c mice orthotopically isografted with the indicated 4T1 mouse breast cancer cells ($1 \times 10^6$), which overexpressed the wild-type mouse $\beta_2$AR (WT $\beta_2$AR) in endogenous $\beta_2$AR knockout 4T1 breast cancer cells (sg$\beta_2$AR-7). A once-weekly intraperitoneal administration of salmeterol xinafoate (2 μg/kg) was initiated from day 7 when the primary tumor reached approximately 100 $mm^3$. The green arrow indicates the time of intraperitoneal injection of salmeterol xinafoate. The primary tumor growth curves were stopped on day 31 when the first dead mouse was observed in the control group (C and D). The colored number is the median survival time (I). The n-values denote the number of mice per group. SX: Salmeterol xinafoate.

(**E**, **F**, and **J**) Average primary tumor growth curves (E), individual primary tumor growth curves (F), and overall survival curves (J) of BALB/c mice orthotopically isografted with the indicated 4T1 mouse breast cancer cells ($1 \times 10^6$), which overexpressed the salmeterol xinafoate-binding-deficient mouse $\beta_2$AR (F193W, F194V, S203T, S204T, H296K, K305D, and Y308F mutations, SBD $\beta_2$AR) in endogenous $\beta_2$AR knockout 4T1 breast cancer cells (sg$\beta_2$AR-7). A once-weekly intraperitoneal administration of salmeterol xinafoate (2 μg/kg) was initiated from day 7 when the primary tumor reached approximately 100 $mm^3$. The green arrow indicates the time of intraperitoneal injection of salmeterol xinafoate. The primary tumor growth curves were stopped on day 31 when the first dead mouse was observed in the control group (E and F). The colored number is the median survival time (J). The n-values denote the number of mice per group. SX: Salmeterol xinafoate.

(**G**, **H**, and **K**) Average primary tumor growth curves (G), individual primary tumor growth curves (H), and overall survival curves (K) of BALB/c mice orthotopically isografted with the indicated 4T1 mouse breast cancer cells ($1 \times 10^6$), which overexpressed the G-protein-binding-deficient mouse $\beta_2$AR (R131A and F139A mutations, GBD $\beta_2$AR) in endogenous $\beta_2$AR knockout 4T1 breast cancer cells (sgβ$_2$AR-7). A once-weekly intraperitoneal administration of salmeterol xinafoate (2 μg/kg) was initiated from day 7 when the primary tumor reached approximately 100 mm$^3$. The green arrow indicates the time of intraperitoneal injection of salmeterol xinafoate. The primary tumor growth curves were stopped on day 31 when the first dead mouse was observed in the control group (G and H). The colored number is the median survival time (K). The n-values denote the number of mice per group. SX: Salmeterol xinafoate.

(**L**, **M**, and **R**) Average primary tumor growth curves (L), individual primary tumor growth curves (M), and overall survival curves (R) of BALB/c mice orthotopically isografted with the indicated 4T1 mouse breast cancer cells ($1 \times 10^6$), which overexpressed the wild-type mouse β$_2$AR (WT β$_2$AR) in endogenous β$_2$AR knockout 4T1 breast cancer cells (sgβ$_2$AR-8). A once-weekly intraperitoneal administration of salmeterol xinafoate (2 μg/kg) was initiated from day 8 when the primary tumor reached approximately 100 mm$^3$. The green arrow indicates the time of intraperitoneal injection of salmeterol xinafoate. The primary tumor growth curves were stopped on day 29 when the first dead mouse was observed in the control group (L and M). The colored number is the median survival time (R). The n-values denote the number of mice per group. SX: Salmeterol xinafoate.

(**N**, **O**, and **S**) Average primary tumor growth curves (N), individual primary tumor growth curves (O), and overall survival curves (S) of BALB/c mice orthotopically isografted with the indicated 4T1 mouse breast cancer cells (1 × $10^6$), which overexpressed the salmeterol xinafoate-binding-deficient mouse $\beta_2$AR (F193W, F194V, S203T, S204T, H296K, K305D, and Y308F mutations, SBD $\beta_2$AR) in endogenous $\beta_2$AR knockout 4T1 breast cancer cells (sg$\beta_2$AR-8). A once-weekly intraperitoneal administration of salmeterol xinafoate (2 μg/kg) was initiated from day 8 when the primary tumor reached approximately 100 $mm^3$. The green arrow indicates the time of intraperitoneal injection of salmeterol xinafoate. The primary tumor growth curves were stopped on day 29 when the first dead mouse was observed in the control group (N and O). The colored number is the median survival time (S). The n-values denote the number of mice per group. SX: Salmeterol xinafoate.

(**P**, **Q**, and **T**) Average primary tumor growth curves (P), individual primary tumor growth curves (Q), and overall survival curves (T) of BALB/c mice orthotopically isografted with the indicated 4T1 mouse breast cancer cells (1 × $10^6$), which overexpressed the G-protein-binding-deficient mouse $\beta_2$AR (R131A and F139A mutations, GBD $\beta_2$AR) in endogenous $\beta_2$AR knockout 4T1 breast cancer cells (sg$\beta_2$AR-8). A once-weekly intraperitoneal administration of salmeterol xinafoate (2 μg/kg) was initiated from day 8 when the primary tumor reached approximately 100 $mm^3$. The green arrow indicates the time of intraperitoneal injection of salmeterol xinafoate. The primary tumor growth curves were stopped on day 29 when the first dead mouse was observed in the control group (P and Q). The colored number is the median survival time (T). The n-values denote the number of mice per group. SX: Salmeterol xinafoate.

Anesthetics were not used in the mouse experiments, except for cancer cells injection, because anesthetics prevented the anticancer effects of salmeterol xinafoate. Mice were fasted for 12 h before salmeterol xinafoate administration. Eight hours after salmeterol xinafoate administration, food was provided. The data are presented as the mean ± s.e.m. values. *P*-values were determined by unpaired two-way ANOVA with uncorrected Fisher's LSD test (C, E, G, L, N, and P) or the log-rank test (I-K and R-T). *$P < 0.05$, ****$P < 0.0001$, and n.s., not significant.

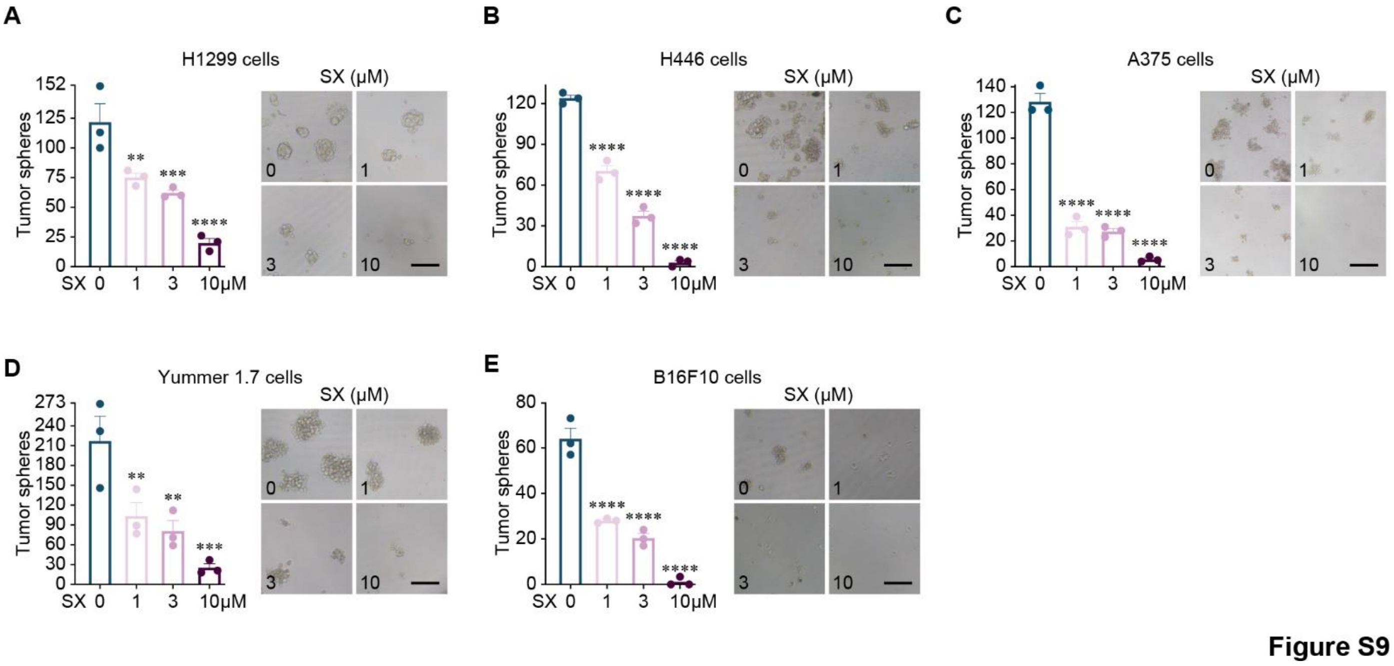


**Figure S9. Salmeterol xinafoate suppresses the capacity of lung cancer and melanoma cells to form tumor spheres**

(**A** and **B**) Number of tumor spheres derived from H1299 human lung cancer cells (1,000, A) and H446 human lung cancer cells (3,000, B) without or with salmeterol xinafoate treatment at the indicated doses (1, 3, or 10 μM) for 7 days (3 independent experiments, left). Representative images of tumor spheres derived from H1299 human lung cancer cells (1,000, A) and H446 human lung cancer cells (3,000, B) without or with salmeterol xinafoate treatment at the indicated doses (1, 3, or 10 μM) for 7 days (right). SX: Salmeterol xinafoate. Scale bar: 100 μm.

(**C-E**) Number of tumor spheres derived from A375 human melanoma cells (1,000, C), Yummer 1.7 mouse melanoma cells (3,000, D), and B16F10 mouse melanoma cells (1,000, E) without or with salmeterol xinafoate treatment at the indicated doses (1, 3, or 10 μM) for 7 days (3 independent experiments, left). Representative images of tumor spheres derived from A375 human melanoma cells (1,000, C), Yummer 1.7 mouse melanoma cells (3,000, D), and B16F10 mouse melanoma cells (1,000, E) without or with salmeterol xinafoate treatment at the indicated doses (1, 3, or 10 μM) for 7 days (right). SX: Salmeterol xinafoate. Scale bar: 100 μm.

The data are presented as the mean ± s.e.m. values. *P*-values were determined by unpaired one-way ANOVA with uncorrected Fisher's LSD test (A-E). $^{**}P < 0.01$, $^{***}P < 0.001$, and $^{****}P < 0.0001$.

**Table S1. Pharmacological characteristics of β2AR agonists**

| Agonist | $pK_D$* | | Functional selectivity ratio β2:β1 | Lipophilic /Hydrophilic (XLOGP3#) | Half-life (hr) | Signaling bias | Type | Refs. |
|---|---|---|---|---|---|---|---|---|
| | β2 | β1 | | | | | | |
| Salmeterol | 9.26±0.06 | 5.73±0.05 | 3388.44 | Lipophilic (3.9) | 5.50 | $G\alpha_s$ | Partial | (*9-14*) |
| Vilanterol | 9.44±0.07 | 6.08±0.07 | 2290.87 | Lipophilic (3.8) | 2.50 | N/A$ | Full | (*15-19*) |
| Formoterol | 7.92±0.03 | 5.86±0.02 | 114.82 | Moderate lipophilic (1.8) | 8.75 | Unbias | Full | (*10, 11, 14, 20-22*) |
| Fenoterol | 6.56±0.02 | 4.90±0.02 | 45.71 | Lipophilic (2.0) | 0.15 | Unbias | Full | (*10, 20, 22-26*) |
| Olodaterol | 8.45±0.02 | 6.80±0.08 | 44.67 | Moderate lipophilic (1.8) | 17.8 | N/A | Full | (*15, 18, 19, 27, 28*) |
| Terbutaline | 5.51±0.04 | 3.90±0.03 | 40.74 | Hydrophilic (0.9) | 3.40 | Unbias | Full | (*10, 22, 23, 29, 30*) |
| Tulobuterol | 6.83±0.09 | 5.62±0.04 | 16.22 | Lipophilic (2.3) | 2.40 | N/A | Partial | (*10, 31-33*) |
| Salbutamol | 5.76±0.03 | 4.74±0.02 | 10.47 | Hydrophilic (0.3) | 4.60 | $G\alpha_s$ | Partial | (*9-11, 14, 20, 24, 34*) |
| Epinephrine | 5.64±0.05 | 4.74±0.04 | 7.94 | Hydrophilic (-1.4) | 0.05 | β-arrestin | Full | (*9-11, 13, 20, 35, 36*) |
| Clenbuterol | 7.44±0.04 | 6.58±0.06 | 7.24 | Hydrophilic (2.2) | 35.0 | Unbias | Partial | (*10, 20, 24, 37-39*) |
| Indacaterol | 7.44±0.08 | 6.81±0.08 | 4.27 | Lipophilic (3.3) | 33.9 | N/A | Full | (*15, 18, 19, 40, 41*) |
| Isoprenaline | 6.64±0.09 | 6.06±0.08 | 3.80 | Hydrophilic (-0.6) | 0.67 | Unbias | Full | (*9, 10, 33, 42, 43*) |
| Norepinephrine | 5.41±0.07 | 5.74±0.03 | 0.47 | Hydrophilic (-1.2) | 0.04 | Unbias | Full | (*10, 22, 44, 45*) |

* The negative logarithm to base 10 of the dissociation constant.
### From PubChem.
$ Not available.

**Table S2. Major binding site on β2AR and the maximum serum concentration of inhaled salmeterol xinafoate or endogenous catecholamines**

| Agonist | Binding site | $C_{max}$ (pg/ml) in plasma | | | Refs. |
|---|---|---|---|---|---|
| | | Cancer | Chronic stress | Acute stress | |
| Salmeterol | H296/K305 | 150* | | | (*9*, *20*, *46*) |
| Epinephrine | S207/F290/ N312 | 49.94±4.10#<br>31.39±4.51$<br>76.94±73.29& | 128.24±12.83 | 43.31±2.39 | (*47*-*51*) |
| Norepinephrine | V114 | 462.03±47.53#<br>74.46±12.52$<br>5753.81±5153.22& | 575.21±52.45 | 586.95±23.92 | (*47*-*52*) |

* Maximum serum concentration in asthmatic patients following a 50 µg dose of inhaled salmeterol powder.
### Patients with oral squamous cell carcinoma.
$ Patients with oropharyngeal squamous cell carcinoma.
& Patients with pheochromocytoma.

**Table S3. Detection of intratumoral salmeterol xinafoate using mass spectrometry***

| Time (min) | Tumor weight (mg) | Average tumor weight (mg) | Intratumoral SX/mouse (pg) | Intratumoral average SX/mouse (pg) | | $C_{max}$ (pg/ml)# (*46*) |
|---|---|---|---|---|---|---|
| 10 | 190.50<br>135.55<br>225.54 | 183.86 ± 26.19 | 305.18<br>666.89<br>220.58 | 397.55 ± 136.86 | | 150 |
| 30 | 150.51<br>130.83<br>256.00 | 179.11 ±38.86 | 175.64<br>89.88<br>328.71 | 198.08 ± 69.85 | | |
| 60 | 304.82<br>141.55<br>232.98 | 226.45 ± 47.25 | 171.01<br>149.90<br>158.66 | 159.86 ± 6.12 | | |
| 120 | 321.48<br>153.55<br>131.66 | 202.23 ± 59.96 | 228.57<br>121.15<br>60.82 | 136.85 ± 49.06 | | |
| 240 | 149.74<br>144.52<br>270.48 | 188.25 ± 41.14 | 43.44<br>24.11<br>129.83 | 65.79 ± 32.50 | | |

* In the MMTV-PyMT mouse model of spontaneous breast cancer (Figure S2L), 7-week-old female mice with established tumors were administered a once-weekly intraperitoneal injection of salmeterol xinafoate (20 μg/kg). The mice were sacrificed at the indicated times (min) after the third injection, and breast tumors (1-3 tumors) were collected. The tumor weight was calculated as the sum of the weight of all tumors per mouse. All tumors from one mouse were mechanically minced together and taken as one sample for mass spectrometry.

### Maximum serum concentration in asthmatic patients following a 50 μg dose of inhaled salmeterol powder.

**Table S4. Comparison between intraperitoneal administration of salmeterol xinafoate in mice and inhalational administration in patients with asthma or chronic obstructive pulmonary disease**

| Mouse dosage (Converted dosage from mouse to human*) | | Clinical dosage (*53*) |
|---|---|---|
| 0.04 µg/mouse/week[#] (5.32 µg/person/week) | | 700 µg/person/week |
| 0.12 µg/mouse/week[$] (15.95 µg/person/week) | | |
| 0.40 µg/mouse/week[&] (53.18 µg/person/week) | | |

* Conversion based on the AUC (area under the concentration-versus-time curve) (*54*) and body surface area (*55*), reference human body weight: 50 kg (*53*), reference mouse body weight: 0.02 kg.
### Intraperitoneal administration of salmeterol xinafoate in mice: 2 µg/kg/week.
$ Intraperitoneal administration of salmeterol xinafoate in mice: 6 µg/kg/week.
& Intraperitoneal administration of salmeterol xinafoate in mice: 20 µg/kg/week.

**Table S5. Comparison between the inhalational administration of salmeterol xinafoate in mice and patients with asthma or chronic obstructive pulmonary disease**

| **Mouse dosage (converted dosage from mouse to humans*)** | | **Clinical dosage** (*53*) | **Toxic dosage** (*53*) |
|---|---|---|---|
| 0.6 μg/mouse/week (122.0 μg/person/week) | | 700 μg/person/week | 10500 μg/person/week |
| 2 μg/mouse/week (406.5 μg/person/week) | | | |
| 6 μg/mouse /week (1219.5 μg/person/week) | | | |

* Conversion based on the body surface area (*55*), reference human body weight: 50 kg (*53*), reference mouse body weight: 0.02 kg.
